# Synovitis in systemic sclerosis is an interferon-driven stromal condition distinct from rheumatoid arthritis

**DOI:** 10.64898/2026.06.30.733140

**Authors:** Celina Geiss, Camino Calvo Cebrian, Miranda Houtman, Ege Ezen, Katerina Apostopoulou, Cristian Iperi, Melpomeni Toitou, Alexandra Khmelevskaya, Marija Lugar, Anne Cauvet, Yannis Djeffal, Mojca Frank Bertoncelj, Sam G. Edalat, Thomas Rauer, Katharina Zachariassen, Kristina Bürki, Cosimo Bruni, Chantal Pauli, Michael Bonelli, Thomas Karonitsch, Yannick Allanore, Raphael Micheroli, Oliver Distler, Caroline Ospelt, Muriel Elhai

## Abstract

Joint involvement is a major driver of disability in systemic sclerosis (SSc), yet its pathophysiology remains poorly understood. In the absence of specific evidence, SSc synovitis is treated by analogy with rheumatoid arthritis (RA). Here, we present the first comprehensive molecular characterization of SSc synovitis, integrating histology, single-cell RNA sequencing, and spatial multi-omics of synovial biopsies from SSc patients, RA patients, and non-inflammatory controls with in vitro validation. We show that SSc synovitis is characterized by distinct pathomechanisms from RA. Histologically, most SSc biopsies displayed a pauci-immune pathotype with sparse immune infiltrates and predominant stromal cells. At molecular level, synovial fibroblasts in SSc were characterized by a disease-specific type I interferon (IFN) response program, in contrast to the TNF-dominant profile of RA, accompanied by dysregulation of the complement cascade. This IFN program extended across multiple synovial cell types, including monocyte-derived macrophages and endothelial cells, and was spatially organized into focal myeloid niches and a diffuse stromal program. Systemically, elevated serum IFN-α2a levels were associated with the presence of clinical synovitis in an independent cohort of SSc patients. We furthermore show that similar IFN-driven programs are shared between skin and synovium in SSc. Genes downregulated by IFNAR1 blockade in SSc skin were enriched in SSc synovium, supporting IFN receptor blockade as a multi-organ target therapeutic strategy. These findings reframe SSc synovitis as a less destructive, IFN-driven stromal condition distinct from RA and provide a mechanistic basis for dedicated clinical trials for joint inflammation in SSc.

**One Sentence Summary:** SSc synovitis is a pauci-immune, IFN-driven stromal condition distinct from RA, supporting IFNAR1 blockade as a therapeutic strategy.

**Graphical abstract:** 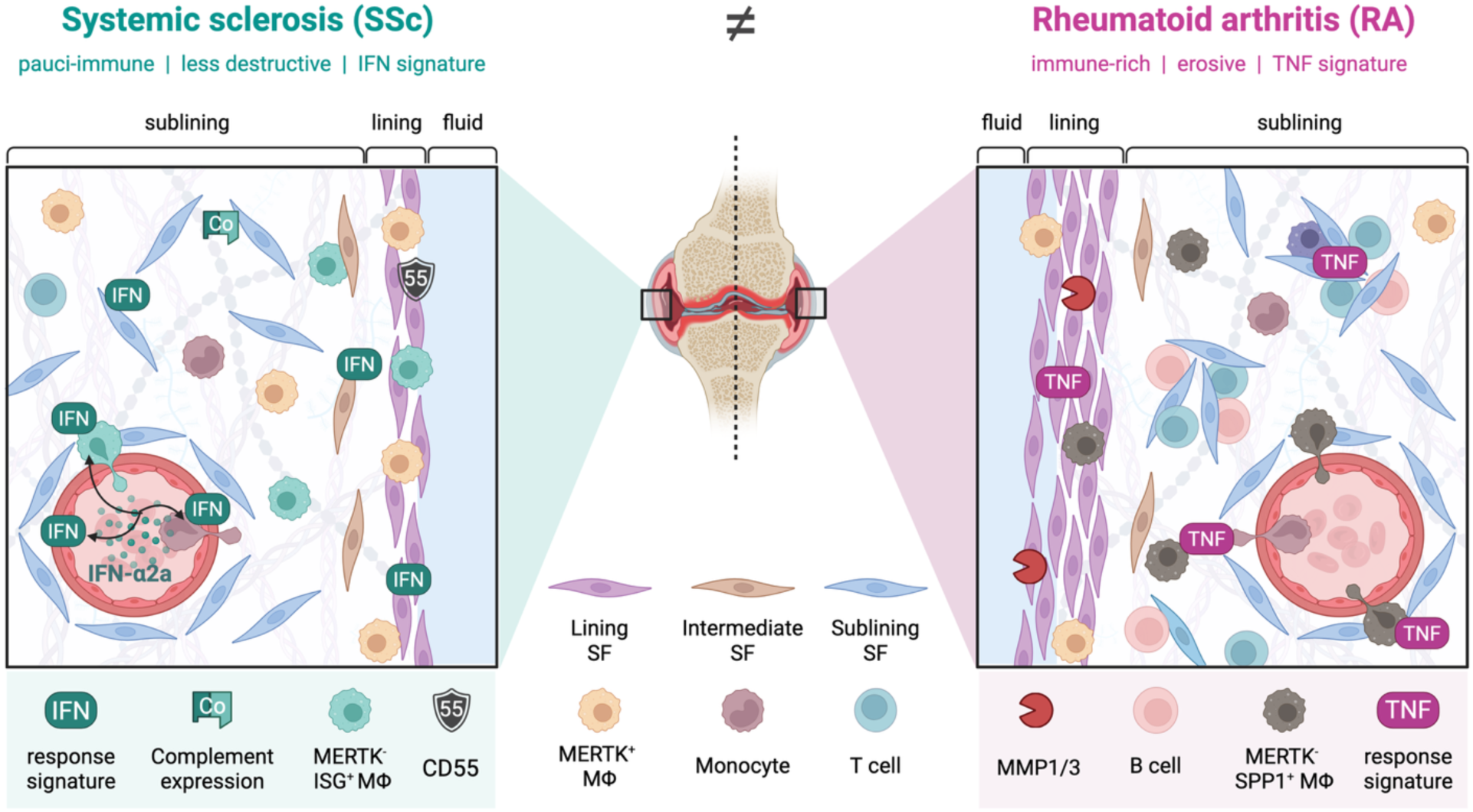

## INTRODUCTION

Systemic sclerosis (SSc) is a rare autoimmune disease, characterized by pathological fibrosis of skin and internal organs. Patients with SSc have high mortality and severely impaired quality of life (*1*). Joint involvement, including synovitis (inflammation of the synovial membrane lining the joints), is a major driver of disability in SSc (*2, 3*). It is a common disease manifestation, affecting up to one third of SSc patients (*4–6*). In addition to its high impact on quality of life, articular involvement is a marker of disease activity and an independent predictor of overall disease progression and death (*5, 7, 8*).

Whether SSc synovitis represents a distinct condition from other forms of inflammatory arthritis is an ongoing debate (*6*). On ultrasound, the pattern of synovitis in SSc differs from that in rheumatoid arthritis (RA), with less inflammation (*9–11*). However, in the absence of evidence-based therapies, SSc patients clinically presenting with joint swelling and pain are treated with immunosuppressive therapies by analogy with RA (*4, 12*). In randomized controlled trials of rituximab, tocilizumab, and abatacept, where SSc joint involvement was assessed as an exploratory or post-hoc measure, no significant effect was shown (*13–15*). A mechanistic understanding of SSc synovitis is therefore needed to determine whether it is pathophysiologically distinct from RA synovitis, which would imply that SSc-specific therapeutic targets need to be identified.

Recent years have revolutionized the understanding of RA pathogenesis by deciphering the heterogeneity of the synovium at the cellular and molecular levels (*16–20*). Nevertheless, the pathogenesis of joint synovitis in SSc remains largely unknown. Only three small studies, published over 50 years ago, showed mild inflammation of the synovium in the early stages of the disease and intense fibrosis in the last stages (*21–23*).

Here, we report the first in-depth molecular characterization of the synovium in SSc using histology, single-cell RNA sequencing (scRNA-seq), spatial multi-omics, and *in vitro* validation (Fig. 1A). Although SSc is considered an immune-mediated disease, we did not find predominant inflammatory infiltrates in the synovium, but a pauci-immune pathotype characterized by the predominance of resident synovial fibroblasts (SFs). Comparison with RA synovium and non-inflammatory control (NIC) synovium revealed hallmark tissue and molecular differences between both arthritides with enrichment in interferon (IFN) signaling pathways across all major synovial cell types in SSc. Systemically elevated IFN levels, shared IFN programs between SSc synovium and skin, and enrichment of the anti-IFNAR1 skin response signature in SSc synovium support the use of IFN-targeted therapies in SSc synovitis.

**Fig. 1.**
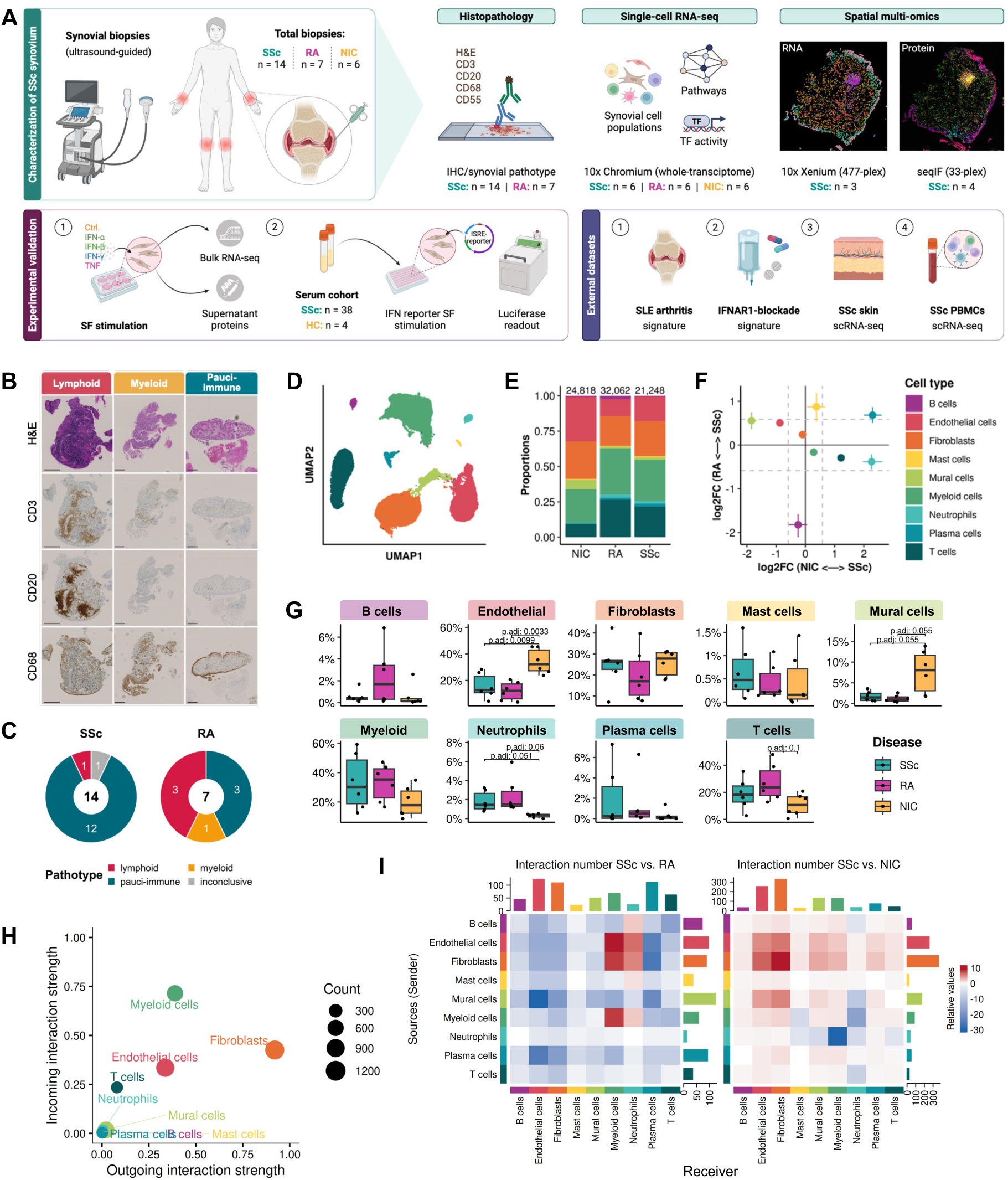
The synovial cellular landscape distinguishes SSc from RA and NIC. (**A**) Schematic study overview: histological pathotype scoring of formalin-fixed paraffin-embedded (FFPE) tissue by immunohistochemistry (IHC), comparative single-cell RNA sequencing (scRNA-seq) of SSc, RA, and NIC (healthy synovium), and spatial profiling of SSc synovium by spatial transcriptomics (10x Xenium) and sequential immunofluorescence (seqIF, Lunaphore COMET), with in vitro synovial fibroblast (SF) stimulation, an ISRE-luciferase serum-stimulation cohort, and integration of external datasets. Created in BioRender. (**B**) Representative images of each synovial pathotype for all biopsies; scale bars, 200 µm. (**C**) Synovial pathotype distribution. (**D**) Integrated Uniform Manifold Approximation and Projection (UMAP) of synovial cell populations in scRNA-seq and (**E**) their proportions per disease group with number above bars. (**F**) Differential cell type abundance (scProportionTest) and (**G**) relative cell type abundances (Wilcoxon rank-sum test with Benjamini-Hochberg (BH) correction). (**H**) Incoming and outgoing cell-cell interaction strength in SSc (CellChat). (**I**) Differential cell-cell interaction numbers; red indicates higher signaling in SSc, blue higher signaling in RA (left) or NIC (right).

## RESULTS

### Clinical and histological characteristics of the patient cohort

A total of 14 patients fulfilling classification criteria for SSc (*24*) with active synovitis requiring treatment were included, presenting with early disease (*25*) and predominantly affected hand and finger joints (*6, 26*), consistent with previously described clinical characteristics of SSc synovitis. For comparison, six seropositive RA patients were included in the study (seven synovial biopsies in total, as one patient was biopsied at both the knee and the wrist, details of clinical characteristics in Table 1). Articular disease was highly active in both groups, as reflected by comparable median DAS28-CRP scores (*27*) (SSc: 4.1; RA: 5.1; Table 1). The synovial biopsy was well tolerated by all patients with no serious side effects, moderate pain and the need for analgesics (NSAIDs, Paracetamol) for one day in four of 20 patients. NIC samples (n=6) with histologically normal synovium were derived from patients undergoing arthroscopy and had no underlying inflammatory joint disease.

**Table 1:** Cohort characteristics.

| Characteristic | SSc (n=14) | RA (n=6) | NIC (n=6) |
| --- | --- | --- | --- |
| Male sex | 3 (21%) | 1 (17%) | 5 (83%) |
| Age (yrs) | 68.0 [61.0–73.0] | 57.5 [54.5–63.5] | 38.0 [29.0–50.8] |
| Disease duration (yrs) | 4.2 [1.2–7.9] | 7.0 [5.4–15.4] |  |
| Antibodies | ACA (n=7), RNA pol III (n=2), SLC70 (n=2), anti-Ku (n=1), isolated ANA (n=2) | Rheumatoid factor (100%), anti-CCP (100%) |  |
| SSc form | limited (n=13), diffuse (n=1) |  |  |
| mRSS | 0.0 [0.0–2.0] |  |  |
| Digital ulcers (no/active/past) | 12/1/1 |  |  |
| Lung fibrosis on HRCT | 7 (50%) |  |  |
| Extensive lung fibrosis | 0 (0%) |  |  |
| Pulmonary arterial hypertension | 1 (7%) |  |  |
| CRP elevation (>5 mg/L) | 3 (21%) | 6 (100%) |  |
| DAS28 ESR | 4.6 [3.9–5.4] | 5.4 [5.0–5.6] (n=5)# |  |
| DAS28 CRP | 4.1 [3.6–4.6] | 5.1 [4.9–5.1] (n=5)# |  |
| Refractory disease | 0 (0%) | 0 (0%) |  |
| Current treatment | bDMARD (n=1) (IL6i)<br>cDMARD (n=1) (LEF)<br>cDMARD (n=1) (LEF+ Plaquenil)<br>cDMARD (n=1) (MPA)<br>cDMARD (n=2) (MTX)<br>cDMARD (n=1) (Plaquenil)<br>naïve (n=5)<br>no (n=2)<br>Steroids (intraarticular and oral) n=0 | bDMARD (n=2) (JAKi)<br>bDMARD (n=1) (TNFi)<br>cDMARD (n=1) (MTX)<br>no (n=2)<br>Steroids (intraarticular and oral) n=0 |  |
| Calcinosis | 2 (14%) |  |  |

Data are median [interquartile range] or n/N (%). ACA: anti-centromere antibodies, ANA: anti-nuclear antibodies. CRP: C reactive protein, DAS28: Disease Activity Score 28-joint count. cDMARD: conventional disease-modifying antirheumatic drug, bDMARD: biological conventional disease-modifying antirheumatic drug, IL6i: IL-6 inhibitor, LEF: leflunomide, HCQ: hydroxychloroquine, MPA: mycophenolate, MTX: methotrexate, TNFi: tumour necrosis factor inhibitor, SZP: salazopyrin, JAKi: janus kinase inhibitor. #data available in 5 patients.

Histologically, SSc synovitis was less inflammatory than RA, with a lower Krenn score (*28*) for all three components (lining layer, stroma and inflammatory infiltrate) and less neutrophil infiltrates (*29*) (Table S1). Notably, there was no significant difference in fibrosis or vascularization between SSc and RA (Table S1). In 12/14 (86%) SSc biopsies, we observed a pauci-immune pathotype (*30*), characterized by sparse immune cells and predominant stromal cells (Fig. 1B). One SSc biopsy was lymphoid and one was inconclusive (mix of lymphoid and myeloid characteristics). RA synovial tissues revealed a heterogeneous tissue composition with balanced pathotypes and pauci-immune synovitis only in 3/7 (43%) synovial biopsies (Fig. 1C) (p = 0.04, Fisher’s exact test, as compared to SSc).

### Synovial fibroblast heterogeneity and abundance in SSc, RA, and NIC

Building on the observed differences in histology, we hypothesized that the molecular profile of SSc synovitis would differ from that of RA. Thus, we generated scRNA-seq data from a subset of biopsies (SSc n = 6, RA n = 6, NIC n = 6, Table S2). We successfully profiled a total of 78,128 cells and identified nine distinct cell type clusters after integration (Fig. 1D and E, Fig. S1A and B). Cell type proportions were broadly consistent with the histological findings, with a tendency towards higher T and B cell abundance in RA (Fig. 1F and G). Cell-cell communication analysis reflected the pauci-immune pathotype, with dominant outgoing signaling from SF in SSc (Fig. 1H, Fig. S1C) compared to RA and NIC and limited immune cell communication (Fig. 1I, Fig. S1D and E).

Given the pauci-immune pathotype observed in SSc patients, characterized by a predominance of stromal cells, and their strong outgoing signaling pattern, we first analyzed SF populations. After re-integration, we obtained a dataset of 15,502 high quality SFs (Fig. 2A, Fig. S2A and B), which segregated into the established *PRG4*^+^ lining (SF1) and multiple sublining SF populations consistent with previous studies (*17, 31*) (Fig. S2C). SF population composition differed across SSc, RA, and NIC. RA samples exhibited a lower relative proportion of *PRG4*^+^ lining SFs compared to SSc and NIC (Fig. 2B) reflecting sublining expansion in RA (*16, 17, 19*) and an enrichment of *CHI3L2*^+^ SF. Notably, *POSTN*^+^ and *FOSB*+ SF were specific to SSc and RA, whereas two SF populations (*APOE*^+^, *FKBP5*^+^) were most abundant in NIC (Fig. 2C and D). Two clusters, *CHI3L2*-expressing SF2 and APOE-expressing SF3 (Fig. 2E and F), were identified as intermediate SF populations displaying transcriptional features of both lining and sublining compartments. The *THY1*^high^ sublining SFs further separated into distinct subsets (Fig. 2E and F, Fig. S2C), including a *COL1A1*^high^ population expressing *POSTN* and *COMP* (SF4), an *FKBP5*+ SF subset (SF5), an *FOSB*^+^*CXCL12*^high^ SF population (SF6), a *CXCL14*^+^*NOTCH3*^+^ SF subset (SF7), and an *MFAP5*^+^*CD34*^+^ (SF8) population resembling a previously described progenitor fibroblast subset (*32*). Overall, no distinct fibroblast population uniquely characteristic of SSc was evident.

**Fig. 2.**
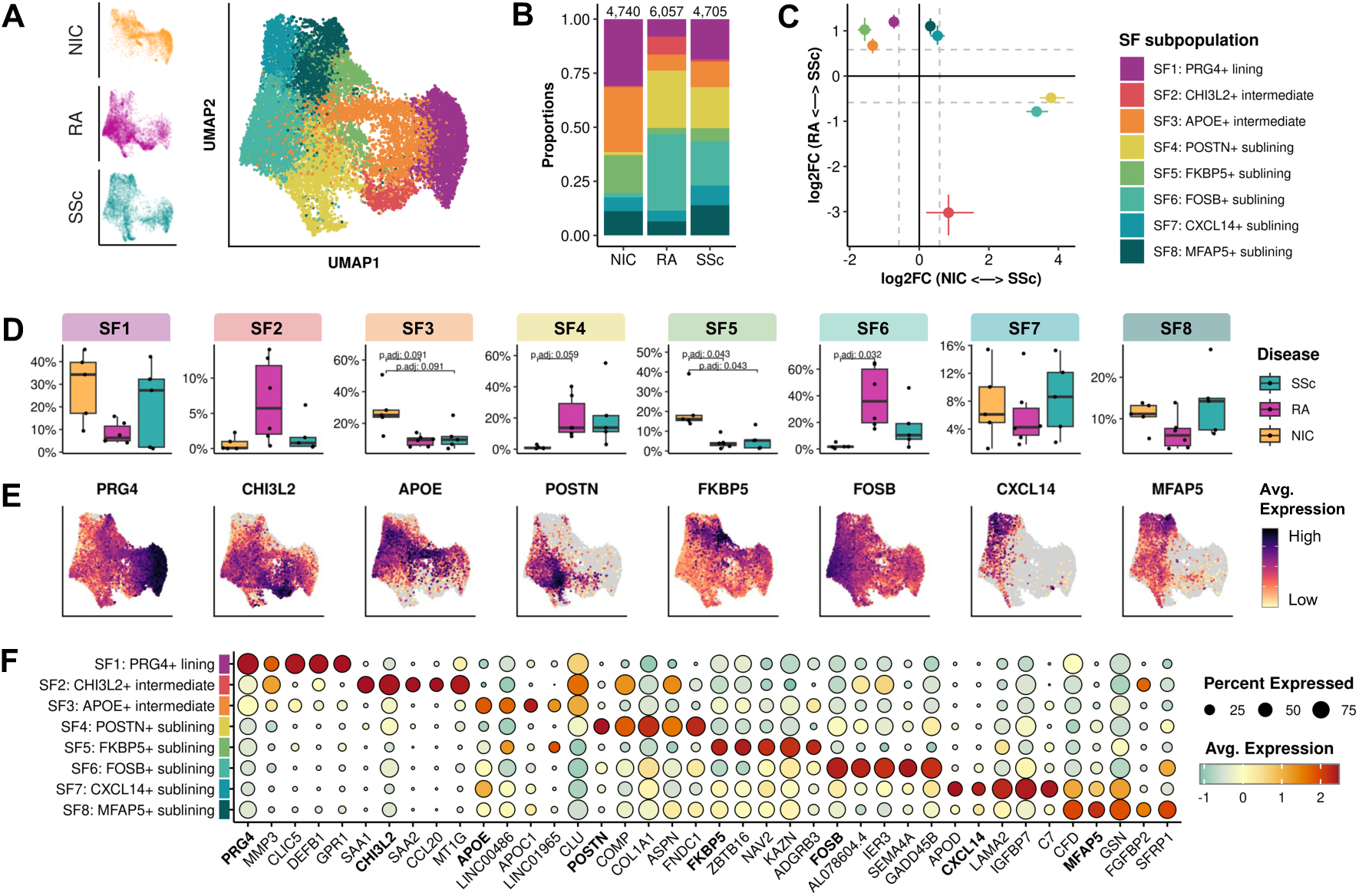
Synovial fibroblast heterogeneity differs between SSc, RA, and NIC. (**A**) UMAP of re-integrated synovial fibroblast (SF) subpopulations. (**B**) Proportions of SF subpopulations per disease group; numbers above bars indicate cells per group. (**C**) Differential abundance of SF subpopulations (scProportionTest). (**D**) Relative SF subpopulation proportions per patient and disease group (Wilcoxon rank-sum test with BH correction). (**E**) Expression of the defining marker gene for each SF subpopulation. (**F**) Top marker gene expression per SF subpopulation; defining marker genes are shown in bold. Dot size indicates the percentage of expressing cells and color the scaled average expression.

### SSc SF transcriptional profile suggests distinct modes of fibroblast activation

We next investigated whether SSc SFs were characterized by disease-specific transcriptional programs rather than distinct population compositions. When comparing to NIC, we found similar pathways enriched in SSc and RA SF (Fig. 3A, Fig. S3A). However, when comparing SSc and RA, several distinct genes and pathways emerged (Fig. 3B). Epithelial mesenchymal transition was enriched in both SSc and RA (Fig. 3B and C, Fig. S3B), but the genes driving this signal differed markedly, suggesting distinct modes of fibroblast activation.

**Fig. 3.**
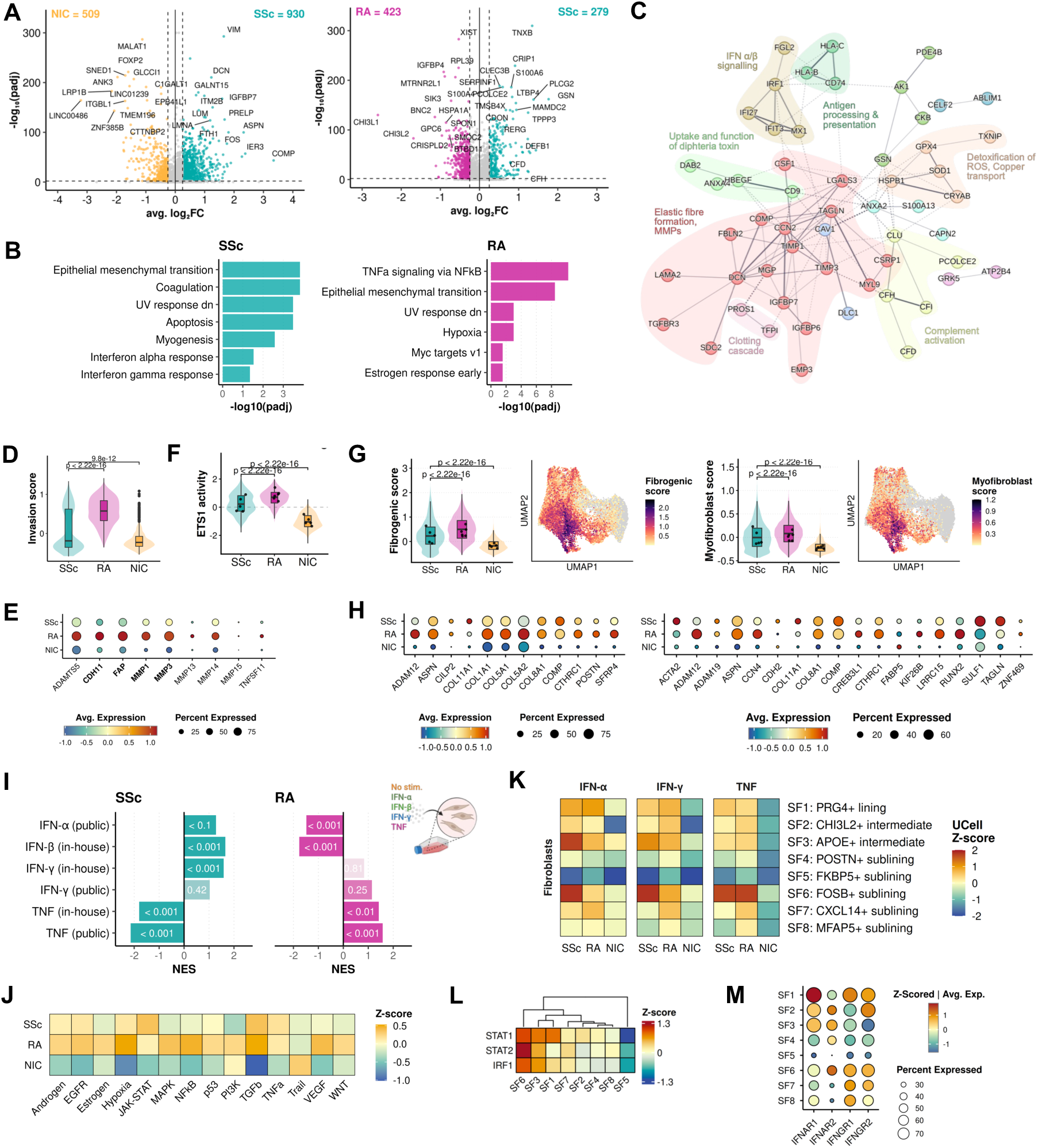
SSc synovial fibroblasts show a distinct transcriptional profile and mode of activation. (**A**) Volcano plots of differentially expressed genes (DEGs) in SSc versus NIC (left) and SSc versus RA (right); numbers indicate up-regulated genes per group. (**B**) Pathway over-representation analysis (ORA) of SSc and RA DEGs (MSigDB Hallmark collection). (**C**) STRING network of the SSc-upregulated genes contributing to the enriched pathways in (B), clustered into functional modules (MCL, inflation = 2). (**D**) Invasion score and (**E**) expression of associated marker genes in SF1; bold genes are significantly differentially expressed between SSc and RA. (**F**) Predicted ETS1 TF activity in SF1 (decoupleR, CollecTRI). (**G**) Synovial fibrogenic and skin myofibroblast scores, dots indicating sample medians, with (**H**) corresponding expression dot plots. (**I**) Gene set enrichment analysis (GSEA) of SSc (left) and RA (right) DEGs against ranked expression changes of in vitro–stimulated SF (in-house and public data). (**J**) PROGENy pathway activity in SF (decoupleR). (**K**) Hallmark IFN-α, IFN-γ, and TNF response gene set scores across SF subpopulations (UCell). (**L**) IFN-associated TF activity across SF subpopulations (decoupleR, CollecTRI). (**M**) IFN receptor expression across SF subpopulations. Comparisons in (**D**), (**F**), (**G**) used the Wilcoxon rank-sum test.

RA SFs were characterized by invasion-associated genes including matrix metalloproteinases and structural collagens. Given that lining SFs are the primary mediators of bone erosion in RA, we examined invasion-associated gene expression (*33*) specifically in the lining compartment. SSc *PRG4*^+^ lining SFs exhibited significantly lower expression of extracellular matrix-degrading markers compared to RA (Fig. 3D and E), and ETS1 transcription factor (TF) activity, linked to invasive fibroblast behavior in RA (*34*), was significantly reduced in SSc (Fig. 3F), indicating that SSc lining SFs lack the invasive program characteristic of RA.

In SSc, SF expressed matrisome-regulatory genes associated with fibrosis, such as *COMP*, *TGFBR3*, and *CCN2*/CTGF (*35, 36*) (Fig. 3C). To assess whether they reflect a profibrotic process in SSc joint, we used two scores, one synovial fibrogenic score derived from RA literature (*37*) and one skin myofibroblast signature (*38*). Both scores were elevated in SSc as well as in RA compared to NIC, with no evidence of SSc-specific enrichment (Fig. 3G and H). Importantly, both scores were also elevated in treatment-naïve SSc and RA (*39*), arguing against a treatment effect (Fig. S3C).

### SSc SFs are characterized by a specific IFN signature

Pathway and network analysis of SSc vs. RA differentially expressed genes (DEGs) in SFs showed specific transcriptional programs. Interferon response (IFN-α and IFN-γ) emerged as prominent signal in SSc, whereas TNF signaling was the most enriched pathway in RA (Fig. 3B and C, Fig. S3D). To independently confirm these findings, we assessed the enrichment of these signatures in *in vitro* IFN and TNF stimulated SFs. The SSc-specific signature was strongly enriched under IFN-α/β stimulation with additional enrichment under IFN-γ, whereas the RA signature was primarily TNF-responsive (Fig. 3I, Fig. S3E). JAK/STAT pathway activity was correspondingly elevated in SSc SFs (Fig. 3J). Across SF populations, IFN-α response enrichment was prominent in the lining (SF1), intermediate (SF2, SF3), and *FOSB*^+^ (SF6) compartments, while the TNF response was elevated in *FOSB*^+^ SF (Fig. 3K). TF activity prediction confirmed elevated STAT1, STAT2, and IRF1 activity in these populations in SSc (Fig. 3L, Fig. S3F). SF populations with IFN-α response signatures also showed the highest *IFNAR1* and *IFNAR2* expression among fibroblast subsets (Fig. 3M).

Consistent with the reduced invasive capacity of lining SF, TNF stimulation induced marked MMP1 and MMP3 accumulation in 3D SF micromasses, whereas IFN stimulation did not elicit a comparable response (Fig. S3G). Together, these findings suggest a disease-specific, broadly distributed IFN program in SSc SF, in contrast to the TNF-dominant profile of RA.

### Dysregulation of complement pathways in SSc SF

In addition to the IFN signature, coagulation and complement cascade pathways were significantly enriched in SSc SF across multiple independent pathway databases (Fig. 3B and C, Fig. S4A and B). Classical complement components and *C3* were upregulated in both diseases compared to NIC (Fig. 4A and B). In the alternative pathway and amplification loop, *CFB* was elevated in both, whereas *CFD*, the rate-limiting enzyme of the alternative pathway, was specifically upregulated in SSc but downregulated in RA, consistent with known reduced *CFD* expression in RA synovium (*40*). Furthermore, multiple complement regulatory factors were increased in SSc SF compared to RA (Fig. 4A and B), indicating simultaneous pathway activation and regulatory control. *CD55*, a known marker for synovial lining (*16, 41*) and central negative regulator of complement activation (*42*), was most strongly expressed in SSc SF, with highest levels in the lining compartment on both transcriptional (SF1, Fig. 4C) and protein levels in the tissue (Fig. 4D). Taken together, SSc SFs exhibit a complex complement dysregulation compared to RA, characterized by both alternative pathway and amplification loop gene expression and upregulation of negative regulatory mechanisms (Fig. S4C).

**Fig. 4.**
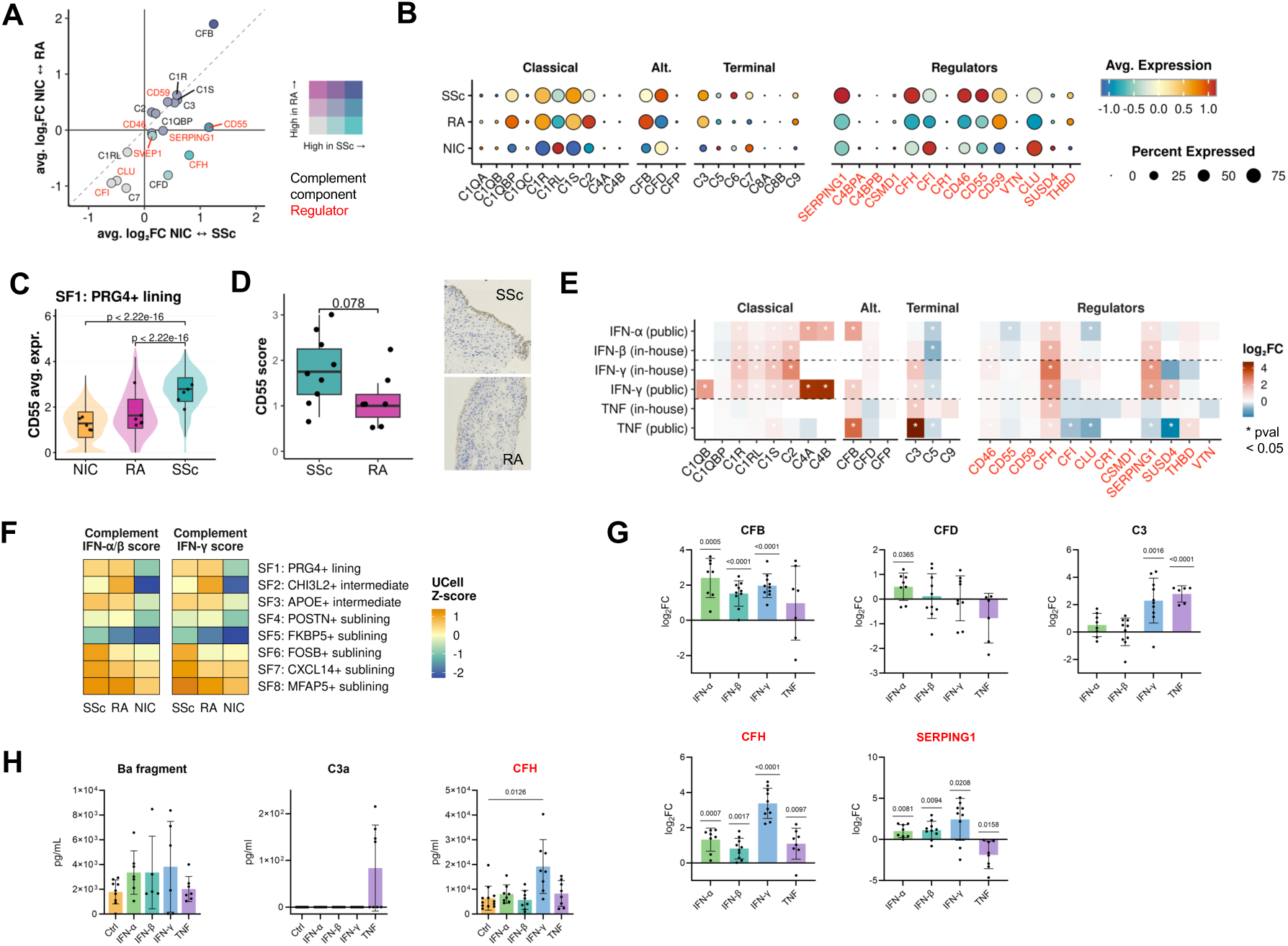
SSc synovial fibroblasts show IFN-driven dysregulation of the complement cascade. (**A**) Differential expression of complement genes (log2FC NIC versus SSc against log2FC NIC versus RA); complement components in black, regulators in red. (**B**) Expression of classical, alternative, and terminal complement pathway genes and complement regulators. (**C**) CD55 expression in SF1; dots indicate sample medians (Wilcoxon rank-sum test). (**D**) CD55 immunohistochemistry in SSc and RA synovial lining (SSc n = 9, RA n = 7); intensity scoring by two blinded observers (ICC = 0.931, 95% CI 0.815–0.974; Wilcoxon rank-sum test). (**E**) Complement gene expression changes after 24 h SF *in vitro* stimulation with type I IFN (IFN-α, IFN-β), type II IFN (IFN-γ), or TNF. (**F**) Module score of IFN-induced complement genes (p < 0.05, log_2_FC > 0 in (E)) across SF subpopulations (UCell). (**G**) qPCR after 48 h of *in vitro* SF stimulation, one-sample t-test (µ=0,; n = 8-10) and (**H**) ELISA of complement components in supernatant: Ba fragment (cleaved CFB; alternative pathway activation), CFH (complement inhibition), and C3a (active C3 convertase); one-way ANOVA with Dunnett’s test versus control. Data points in (G) and (H) represent individual biological replicates; bars indicate mean ± SD.

### IFN stimulation induces SSc-specific complement expression pattern in SF

Given the co-occurrence of IFN and complement signatures in SSc, we investigated whether IFN directly regulates complement expression in SFs. While IFN-complement crosstalk has been described in systemic lupus erythematosus (SLE) (*43, 44*), its role in the synovium and in SSc remains poorly understood (*40, 42*). Revisiting the *in vitro* stimulation transcriptomics, IFN-α/β and IFN-γ upregulated classical complement components and regulators, closely resembling the SSc SF pattern, whereas TNF stimulation induced only a partial response, mirroring the RA milieu (Fig. 4E). Based on these IFN-induced complement genes, SF7 and SF8 emerged as the primary complement-associated populations, with weaker enrichment in SF6 (Fig. 4F). qPCR independently confirmed concurrent upregulation of complement components and regulators upon IFN stimulation of SF *in vitro* (Fig. 4G, Fig. S5). At the protein level, IFN stimulation increased the Ba fragment (cleaved CFB), indicating alternative pathway activation, but also the alternative pathway inhibitor CFH, whereas C3a showed no consistent induction (Fig. 4H). Together, IFN stimulation may drive a complement program that recapitulates the pattern observed in SSc SFs.

### IFN response across multiple synovial cell types in SSc

Having established an IFN program in SF, we next tested whether this signature extends to other synovial cell types. Venous and arterial endothelial cells displayed an elevated IFN-α response signature in SSc compared to NIC, while this was not observed in RA (Fig. 5A, Fig. S6).

**Fig. 5.**
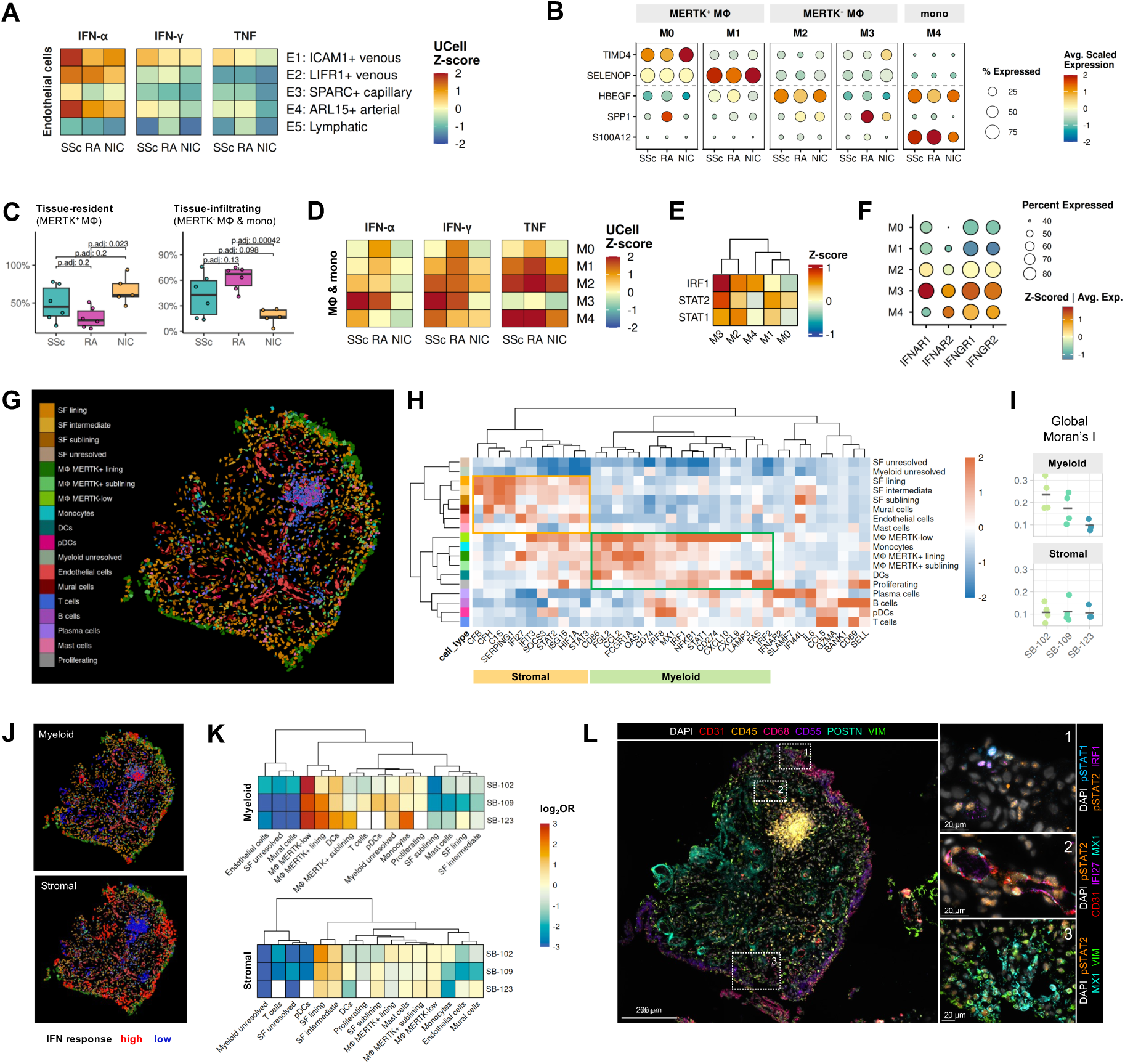
The IFN response spans stromal and myeloid compartments and is spatially organized in SSc synovium. (**A**) Hallmark IFN-α, IFN-γ, and TNF response gene set scores across endothelial cell populations (UCell). (**B**) Marker gene expression for macrophage (MΦ) populations and monocytes. (**C**) Proportions of tissue-resident populations and tissue-infiltrating myeloid populations (Wilcoxon rank-sum test, BH correction). (**D**) Hallmark IFN-α, IFN-γ, and TNF response gene set scores across MΦ and monocytes (UCell). (**E**) IFN-associated TF activity (decoupleR, CollecTRI) and (**F**) IFN receptor expression across MΦ and monocyte populations in SSc. (**G**) Cell type annotation in Xenium spatial transcriptomics (n = 3), representative section (patient SB-109). (**H**) Hallmark IFN-α/IFN-γ response genes within the Xenium panel across cell types; hierarchical clustering (without spatial information) resolves a stromal and a myeloid IFN program. (**I**) Spatial autocorrelation (global Moran’s I) of myeloid and stromal IFN response programs; dots indicate tissue pieces, bars patient means. (**J**) Spatial distribution of the myeloid and stromal IFN response niches (local Moran’s I based on UCell). (**K**) Cell type enrichment (log_2_ odds ratio, OR) in spatially defined IFN-high niches; white indicates absence of cell type. (**L**) Representative seqIF of consecutive section to (G) showing myeloid (1), endothelial (2), and stromal (3) niches.

T cells showed few differences and no enrichment of IFN response pathways between SSc, RA, or NIC. UCell scoring indicated some IFN response in NK cells and regulatory T cells; however, these populations were small in SSc, precluding robust conclusions (Fig. S7).

Furthermore, we characterized four populations of macrophages (M0: *MERTK*^+^*TIMD4*^+^, M1: *MERTK*^+^*SELENOP*^+^, M2: *MERTK*^-^*HBEGF*^+^, M3: *MERTK*^-^*SPP1*^+^) and one of monocytes (M4: *S100A12*^+^) in line with literature (*17, 45, 46*) (Fig. 5B, Fig. S8). Consistent with the homeostatic composition of healthy synovium, NIC was dominated by tissue-resident *MERTK*⁺ macrophages M0 and M1 (Fig. 5C). Tissue-infiltrating, monocyte-derived *MERTK*^-^ populations (M2, M3), and M4 monocytes were present in both, SSc and RA, with a higher abundance in RA. These populations recapitulated features previously described in RA, however they also showed disease-specific differences. The M2 population shared features with the previously described *HBEGF*^+^ inflammatory macrophage state (*46*), but lacked characteristic *EREG* expression in SSc (Fig. S8E). The M3 population was defined by SPP1 as its top marker gene, however *SPP1* expression was restricted to RA (Fig. 5B). Since both, M2 and M3, have been linked to joint invasion in RA (*16, 45, 47*), these differences might contribute to less synovial invasion observed in SSc.

In RA, tissue-resident *MERTK*^+^ macrophages (M0 and M1) showed strong TNF and IFN response signature, suggesting local inflammatory activation (Fig. 5D). In contrast, in SSc the IFN response signature was restricted to monocytes and monocyte-derived macrophage populations. TF activity prediction confirmed strong enrichment of IFN-associated transcription factors in these *MERTK*⁻ subsets (Fig. 5E), corroborated by elevated IFN receptor expression in the same populations (Fig. 5F).

To test whether the IFN response in these *MERTK*⁻ populations reflects systemic priming prior to synovial infiltration we examined circulating monocytes from an independent publicly available PBMC dataset (*48*) and observed that both classical and non-classical monocytes from SSc patients showed elevated IFN-α and IFN-γ response scores compared to healthy controls (Fig. S9). Together, this pattern supports a model in which monocyte-derived populations infiltrating SSc synovium are primed by systemic IFN rather than activated locally.

### Spatial organization of the synovial IFN response

To resolve the tissue architecture of the synovial IFN program, we analyzed all biopsies from SSc patients with sufficient available FFPE tissue by single-cell spatial transcriptomics (n = 3, Fig. 5G, Fig. S10) and sequential immunofluorescence (seqIF, n = 4). Clustering of IFN response gene expression (IFN-α and IFN-γ) across cell types identified two transcriptionally distinct programs: a stromal program and a myeloid program (Fig. 5H). Spatial autocorrelation analysis (global Moran’s I) confirmed significant clustering of both programs across all tissue pieces, with the myeloid program showing higher autocorrelation than the stromal program (Fig. 5I) indicating stronger spatial organization of the myeloid IFN program. LISA-based (local Moran’s I) niche classification identified discrete myeloid niches enriched for *MERTK*⁻ macrophages, monocytes, and dendritic cells, spatially segregated from stromal populations that were enriched in lining and intermediate SF (Fig. 5J and K).

These spatially defined niches were independently validated at the protein level by seqIF on consecutive tissue sections. Nuclear co-localization of pSTAT1 and IRF1 was confined to myeloid-rich zones (Fig. 5L), indicating focal JAK/STAT transcription factor activity in myeloid niches. In contrast, pSTAT2 showed diffuse nuclear localization across the lining zone with additional detection in endothelial cells, corroborated by MX1 expression in the same compartment (Fig. 5L). Together, these findings establish a dual spatial architecture of the SSc synovial IFN response: discrete, JAK/STAT-active myeloid niches embedded within a broader, diffuse stromal program that extends to the synovial vasculature.

### Systemic IFN-α2a levels are associated with synovitis in SSc

To see whether this widespread IFN response signature has a local origin, we investigated known sources of IFN within the synovium. No type I IFN transcripts were detected, and *IFNG* expression was restricted to T cells without enrichment in SSc (Fig. S11A). Local cGAS-STING-driven IFN-β synthesis appeared unlikely given low predicted IRF3 activity in fibroblasts and myeloid populations (*49*) (Fig. S11B). Plasmacytoid dendritic cells (pDCs), established producers of type I IFN in SSc (*50*), were detectable in spatial data but present at low abundance and showed no consistent enrichment in IFN response-high niches across patients (Fig. 5K), arguing against local pDC-driven IFN production as the primary driver. Although the absence of detectable *IFN* transcripts does not exclude local production, the prominent IFN response signature in cell populations with direct exposure to systemic circulation – monocyte-derived macrophages, monocytes, and endothelial cells – together with reports of elevated systemic type I IFN activity in SSc (*51–53*), prompted us to investigate external sources as potential drivers of the synovial IFN response signature.

To explore whether elevated systemic IFN levels are associated with joint involvement in SSc, we analyzed an independent cohort of 132 SSc patients derived from (*54*) without overlap disease (Table S3). In a binary logistic regression, higher IFN-α2a levels were significantly associated with the presence of synovitis (OR = 3.74, p = 0.013; AUC = 0.65; Fig. 6A). To assess specificity, we performed a multinomial logistic regression with no joint involvement as reference. IFN-α2a levels were elevated specifically in patients with synovitis (β = 1.25, OR ≈ 3.5, p = 0.019), but not in those with tender joints only (β = –0.65, p = 0.21; Fig. 6B), indicating that the association is specific to objective joint inflammation rather than pain-related symptoms alone.

**Fig. 6.**
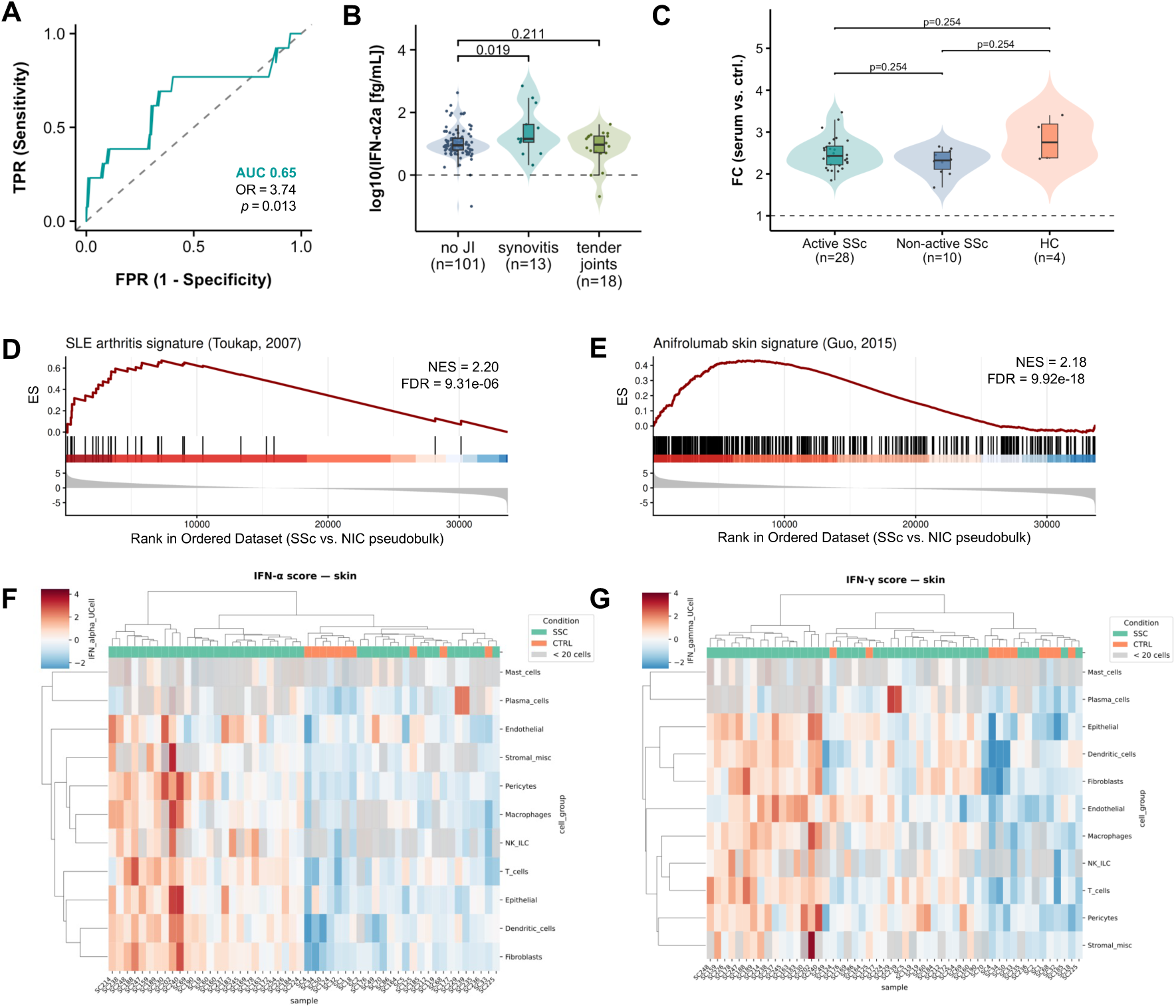
Systemic IFN activity and therapeutic implications. (**A**) ROC curve of a binary logistic regression model predicting the presence of synovitis from serum IFN-α2a levels in an independent cohort of 132 SSc patients. (**B**) Serum IFN-α2a levels of SSc patients in (A) stratified by joint phenotype. Dots represent individual patients; boxes show median and IQR. P-values are derived from multinomial logistic regression with no joint involvement as reference. (**C**) ISRE-luciferase reporter activation in SF stimulated with sera from SSc patients (n = 38) grouped by disease activity or healthy controls (HC, n = 4); Wilcoxon rank-sum test with BH correction. (**D**) GSEA of a SLE arthritis gene signature and (**E**) of genes downregulated by IFNAR1 blockade with anifrolumab in SSc skin against SSc vs. NIC pseudobulk DEGs ranked by log₂FC. (**F**) IFN-α and (**G**) IFN-γ response UCell Z-scores across cell types in publicly available SSc skin datasets; Z-scores computed per cell type relative to healthy controls (CTRL).

To directly test whether circulating IFN concentrations are sufficient to induce an IFN response signature in SF *in vitro*, we stimulated two IFN-stimulated response elements (ISRE)-luciferase reporter cell lines with a second independent cohort of SSc patient sera (n = 38, Table S4). Serum stimulation activated the reporter system, but SSc patient sera did not induce significantly higher reporter activity compared to healthy controls (Fig. 6C, Fig. S11C), suggesting that circulating IFN concentrations alone may be insufficient to fully account for the synovial IFN response signature observed in SSc SF.

### IFN type I signaling in SSc synovitis as a candidate therapeutic target

The consistent type I IFN response signature across synovial cell types, supported by its association with systemic IFN activity, prompted us to ask whether this program may be therapeutically targetable. Type I IFN targeted therapies have shown efficacy in SLE arthritis (*55–57*) and in SSc skin (*58*). Pseudobulk GSEA revealed strong enrichment of a SLE arthritis signature (*59*) in SSc synovium compared to NIC (Fig. 6D), indicating shared transcriptional features. Consistent with this, pseudobulk GSEA revealed significant enrichment of genes downregulated following IFNAR1 blockade with anifrolumab in SSc skin (*58*) among SSc synovial DEGs (Fig. 6E). Furthermore, SSc synovium and skin shared common IFN programs (Fig. 6F and G). Together, these findings support IFNAR1 blockade as a rational therapeutic strategy in SSc synovitis, warranting evaluation in dedicated clinical studies.

## DISCUSSION

SSc synovitis is often managed by analogy with other inflammatory arthritides like RA (*60, 61*), in the absence of mechanistic understanding of the underlying biology. By integrating histology, single-cell and spatial omics, and *in vitro* functional assays, we provide a molecular framework for SSc synovitis that reveals it is not a milder or attenuated form of RA inflammation, but a biologically distinct condition defined by a pauci-immune, IFN-dominant stromal program – a distinction with direct implications for target selection in SSc joint disease.

The pauci-immune pathotype observed in the majority of SSc biopsies contrasts with the inflammatory infiltrates characteristic of RA. The absence of SPP1^+^ macrophages, previously linked to synovial invasion and bone erosion in RA (*47*), and the reduced invasive capacity signature of lining SF, evidenced by lower ETS1 activity (*34*) and reduced expression of matrix-degrading enzymes, suggest that the SSc synovium lacks the destructive joint remodeling machinery seen in RA (*33*). This may explain the lower frequency and different pattern of erosions in SSc synovitis (*6, 26, 62*).

In contrast to the TNF-dominant profile of RA, TNF pathway activity in SSc synovium was only modestly elevated, and TNF blockade is not routinely recommended in SSc given concerns regarding fibrosis progression and the absence of controlled trial evidence (*63*). Instead, we show that the specific transcriptional feature of SSc synovitis is not inflammation in the classical sense, but a coordinated IFN response program. This is biologically plausible, as type I IFNs are established drivers of SSc pathogenesis in skin and lung (*64*). Our data extend this paradigm to the joint, where the IFN program is distributed across multiple SF populations and infiltrating *MERTK*^-^macrophages and monocytes, supported by elevated JAK/STAT pathway activity and enrichment of major IFN-response TFs at both the activity prediction and protein level. Spatially, this program is structured into discrete myeloid niches alongside a more diffuse stromal enrichment, establishing a multi-compartment IFN program as the molecular hallmark of SSc synovitis.

Although both type I and type II IFN response signatures are detectable in SSc synovium, their substantial transcriptional overlap makes definitive discrimination challenging. *IFNG* expression was restricted to T cells and not enriched in SSc, and T cells showed no consistent enrichment in IFN response-high niches in spatial data, together supporting a predominant type I IFN contribution. The association of serum IFN-α2a with clinical synovitis, together with preferential IFN response enrichment in monocytes and monocyte-derived macrophages and elevated IFN response scores in circulating monocytes from SSc patients, supports a systemic contribution to the myeloid IFN signature. However, serum stimulation was insufficient to fully account for the stromal IFN response. SSc patient sera did not activate an ISRE-luciferase reporter system above healthy control levels, raising the possibility of additional local IFN production within the joint. Despite interrogating multiple candidate mechanisms, the cellular source of this putative local response remains unresolved.

Beyond canonical ISG induction, SSc SFs show concurrent dysregulation of the complement cascade, with simultaneous upregulation of alternative pathway components and complement regulatory factors on both the transcriptional and protein levels – suggesting a state of pathway activation alongside a regulatory counter-response. IFN-complement crosstalk has also been described in SLE, where complement consumption and arthritis was linked to IFN-dependent mechanisms (*43, 44*). Further work using in-situ complementomics will be needed to determine whether the alternative pathway is functionally active in SSc synovial tissue (*65*).

A number of limitations in this study should be acknowledged. The rarity of SSc synovial biopsies restricted cohort size, and replication in larger, prospectively collected cohorts is needed to confirm the generalizability of our findings across the full spectrum of SSc synovitis. The NIC cohort, while histologically normal, was demographically skewed towards younger male patients, which may limit direct comparability with the predominantly older, female SSc cohort. Due to limited availability of sample material, spatial transcriptomics was performed in only three patients, and larger datasets will be required to fully characterize the architecture of IFN niches in SSc synovium. Our anifrolumab enrichment analysis is derived from a small open-label Phase I skin trial (*58*), and its relevance to joint disease requires prospective validation.

Several strengths of this study warrant emphasis: It provides the first comprehensive molecular characterization of SSc synovitis, addressing a critical gap that has persisted for over five decades. Synovial biopsies in SSc are technically demanding and carry procedural risks in a population with impaired wound healing, rendering this cohort exceptionally valuable. Beyond joint disease, this study establishes a mechanistic framework linking systemic IFN activity to tissue-level programs across multiple organs in SSc, advancing the conceptual understanding of IFN as a multi-organ pathogenic driver.

Altogether, these findings provide a mechanistic rationale for targeting IFN signaling in SSc synovitis and for including joint outcomes in future studies evaluating IFN-targeted therapies in SSc. The transcriptional similarity between SSc synovium and SLE arthritis (*59*) suggests shared IFN-driven pathomechanisms at the joint level. In SLE, IFN blockade reduced joint inflammation (*57*), providing proof-of-concept that synovitis in an IFN-high disease context is amenable to IFN-targeted therapy. The absence of a dominant SSc-specific fibrogenic program in our early cohort suggests a therapeutic window before joint contractures and irreversible fibrosis occur (*6, 21–23, 66*). Anifrolumab is currently being evaluated in SSc for skin and lung involvement (*67*), and the evidence presented here provides a compelling rationale for conducting a trial in SSc synovitis.

Taken together, these findings have the potential to inform a paradigm shift from empirical, RA-analogous treatment towards evidence-based therapeutic strategies in SSc synovitis. By reframing SSc synovitis as an IFN-driven stromal condition distinct from RA, they open a new avenue for targeted therapy in a disease where joint involvement represents a long-standing unmet need.

## MATERIALS AND METHODS

### Study Design

The objective of this study was to characterize the cellular and molecular composition of the synovial microenvironment in SSc and to compare it with RA and non-inflammatory controls (NIC). Synovial biopsies were obtained from consecutive SSc patients fulfilling the 2013 ACR/EULAR classification criteria with active synovitis confirmed by ultrasound, treated at the rheumatology department of the University Hospital Zurich between 09/2020 and 08/2025. As inflammatory controls, synovial biopsies from seropositive RA patients fulfilling the 2010 ACR/EULAR classification criteria were included. NIC samples were obtained from patients undergoing arthroscopy with histologically normal synovium. Patients with SSc overlap syndromes or positive RF or anti-CCP were excluded to ensure diagnostic clarity. CD55 immunohistochemistry was scored independently by two blinded observers. Spatial organization of the synovial IFN program was assessed by single-cell spatial transcriptomics (10x Genomics Xenium, 477-plex) in three SSc patients with available material and validated at the protein level by seqIF (Lunaphore COMET, 33-plex) on consecutive FFPE sections. To investigate the functional consequences of IFN signaling, synovial fibroblasts derived from non-inflammatory donors were stimulated with recombinant cytokines in 2D culture and 3D micromass models. The capacity of circulating SSc IFN to activate fibroblasts was assessed using ISRE-luciferase reporter cell lines stimulated with SSc patient sera; details are provided in the Supplementary Materials. The association between systemic IFN activity and clinical synovitis was examined in an independent, previously published cohort (*54*) of 132 SSc patients. The study was approved by the Ethics Committee of the Canton of Zurich (BASEC-Nr: 2019-00674, 2021-00092, 2021-01818). All participants provided written informed consent.

### Statistical analysis

All statistical analyses were performed in R. scRNA-seq data were corrected for ambient RNA (SoupX) and doublets (scDblFinder), analyzed in Seurat, using Harmony for integration. Cell populations were annotated manually based on prior knowledge and re-integrated with STACAS for subpopulation identification. Differential cell type abundance was assessed using Pochi and scProportionTest. Pairwise differential expression analysis was performed using MAST, including biopsy as a latent variable, with Benjamini-Hochberg correction, with significance defined at BH-adjusted p < 0.01 and |log2FC| > 0.25. Over-representation analysis was performed using clusterProfiler, gene set scores were calculated using UCell. Pathway and transcription factor activity were estimated using decoupleR. Spatial cell type annotations were performed manually per patient and subpopulations inferred with RCTD (spacexr) using the SSc scRNA-seq data as a reference. Spatial autocorrelation of MSigDB Hallmark IFN response UCell scores was assessed using global and local Moran’s I on a k-nearest neighbor spatial weight matrix (k = 15). Cell type enrichment in spatially defined IFN-high versus IFN-low niches was quantified as log2 odds ratio per patient. Histological pathotype proportions were compared using Fisher’s exact test. Association of synovitis with serum IFN-α2a concentrations was assessed by binary logistic regression, with discriminatory performance quantified by area under the receiver operating characteristic curve (AUC). For *in vitro* experiments, qPCR and ELISA data were analyzed using GraphPad Prism; statistical tests are indicated in the respective figure legends. For all analyses, a two-sided p-value < 0.05 was considered statistically significant unless otherwise stated.

## Acknowledgments

We thank all patients for their contribution. We thank our patient research partners A. Eisenring and J. Messmer for ensuring the study is targeting patient’s needs. We thank P. Künzler, A. Laimbacher, S. Dettwiler and F. Prutek for their excellent technical assistance. We thank N. Schneider and P. Huber for administrative support. We thank P. Rossbach and M. Marks for sample collection. We thank the C. Urban, P. Guegen and the Functional Genomics Center Zurich for conducting the transcriptomics data generation and for technical support. We thank O. Hanley, N. Schilling and the Center for Microscopy and Image Analysis (ZMB) at the University of Zurich for supporting the spatial proteomics data generation. We thank

## Funding

Foundation for Research in Rheumatology (FOREUM), Project: “Deciphering synovitis in systemic sclerosis”, ME

Novartis Foundation for Medical-Biological Research, Application #23C170, Project: “Deciphering synovitis in systemic sclerosis”, ME

UZH Advanced Clinician Scientist Program, University of Zurich, Project: “Deciphering arthritis in systemic sclerosis and pauci-immune arthritis “, ME

Filling the Gap, University of Zurich, Project: “Deciperhing the pauci-immune synovitis in joint diseases”, ME

Association des sclérodermiques de France, Project: „Charactérisation de la synovite dans la sclérodermie systémique“, ME

Theodor und Ida Herzog-Egli Stiftung, Project: „Deciphering synovial fibrosis in joint diseases“, ME

10x Genomics & Functional Genomics Center Zurich Core Lab Grant; Project: “A Spatial Map of Arthritis in Systemic Sclerosis“, CG

EULAR, Voucher ID Q125RSV232, Project: “Mulitplex protein analysis of synovium”, ML

R’Equip, Swiss National Science Foundation, Application: 316030_221562, Project: “Ultrahigh content imaging microscopy”, CO

## Author contributions

Conceptualization: CG, ME, CO, OD

Data curation: CG, CCC, EE, CI, CB, AC, YA

Formal analysis: CG, CCC, EE, CI, MH, YD, CP

Funding acquisition: ME, CO, CG, ML, OD

Investigation: CG, CCC, MT, EE, MH, AK, KA, CP, RM, KZ, CP, TR, MFB, SGE

Methodology: CG, MH, CCC, MT, EE, AK, KA, ML, MFB, SGE

Project administration: ME, CO, CG, OD

Resources: KB, CB, KA, TK, MB, AC, YA, TR

Supervision: ME, CO, OD

Validation: CG, CCC, EE, MT, CI

Visualization: CG, CCC, EE, CI

Writing – original draft: CG, ME, CO, CCC, EE, CI

Writing – review & editing: CG, ME, CO, OD, EE, CI, CB, CCC, ML, YD

All authors reviewed and approved the final manuscript.

