## Supplementary Material (Methods, Figures S1-S11) for "Synovitis in systemic sclerosis is an interferon-driven stromal condition distinct from rheumatoid arthritis"

Celina Geiss *et al.*

**The PDF file includes :**

Materials & Methods

Figures S1 to S11

References (*68–110*)

**Other Supplementary Material for this manuscript includes the following:**

Tables S1 to S6

### MATERIALS & METHODS

#### Study design and patients

The study took place in our rheumatology department in Zurich, Switzerland between September 2020 and August 2025. One biopsy performed in 2015 was reanalyzed in 2022 for histology (no scRNA-sequencing analysis performed). The study included consecutive SSc patients responding to the 2013 American College of Rheumatology (ACR)/European League against Rheumatism (EULAR) criteria for SSc *(24)* with active synovitis assessed by ultrasound. SSc patients with overlap with another autoimmune rheumatic disease *(68)* or positive for RF (rheumatoid factor) or anti-CCP (cyclic citrullinated peptide) were not included. We also included patients from our cohort of synovial biopsies from RA patients *(31)* responding to the 2010 ACR/EULAR classification criteria for RA *(69)*, which were positive for RF and anti-CCP, to ensure a homogeneous control group and avoid misdiagnosis of RA *(70)*. Samples from arthroscopy patients without inflammatory joint disease and histologically normal synovium and were designated as non-inflammatory controls (NIC).

The following characteristics were collected: age, sex, disease duration (defined by the first non-Raynaud symptom in SSc), antibodies, cutaneous form according to LeRoy *(71)*, presence of digital ulcers, presence and extent of lung fibrosis on high-resolution computed tomography (HRCT), extensive lung disease according to *(72)*, pulmonary arterial hypertension according to right heart catheterisation *(73)* and CRP levels (mg/L). CRP elevation was defined by CRP > 5 mg/L. Articular disease was assessed using the Disease Activity Score 28-joint count (DAS28-ESR/CRP) *(74)*. Erosion and calcinosis were assessed on X-rays of both hands *(75)*. Treatment was collected. A refractory disease was considered in case of failure to multiple conventional disease-modifying antirheumatic drugs (csDMARDs) and biological DMARDs (bDMARDs) *(76)*.

Two trained rheumatologists performed ultrasound at the joint according to Outcome Measures in Rheumatology in Clinical Trials (OMERACT) Ultrasound Task Force *(77)*. In case of synovitis of at least grade 1, ultrasound-guided synovial biopsy was performed for further diagnosis (8 wrists, 2 MCP and 1 knee) by two experienced rheumatologists (RM; KZ). 15 to 20 separate fragments from different sites in the same joint were collected. Half were used for histological assessment, half for scRNA-sequencing. Exclusion criteria for biopsy were as follows: severe vasculopathy with previous or current digital ulcers for the metacarpophalangeal joints (MCP) in SSc, Quick < 65%, INR > 1.3 or platelet count < 100'000/μL. Due to the risk of impaired healing in SSc, no biopsy was performed on PIP (proximal interphalangeal) joints in SSc patients.

**Histological assessment of synovial tissue and quality control**

Synovial tissue quality and grading of synovitis were evaluated in formalin-fixed, paraffin-embedded sections by histologic analysis (H&E staining) by a pathologist (CP). Specimens were identified as synovium by the presence of a lining layer. Samples consisting of dense fibrous tissue, joint capsule or other tissues were determined not to be synovium. For each histological assessment, we generated pooled data from 8–10 separate fragments from different sites in the same joint. Thus, this should be representative of the whole tissue and mitigate much of the biopsy site-to-site variability. Synovitis score was assessed by evaluation of the thickness of the lining cell layer, the cellular density of synovial stroma and leukocyte infiltration as described by Krenn *et al*. *(78)*. Vascularization was assessed by counting the number of CD31+ cells (Abcam, ab28364) in five consecutive images at 20× magnification. Synovial tissue was stained with CD3 (Abcam, ab16669), CD20 (Agilent, M075501-2), CD68 (Agilent, M081401), and CD138 (Agilent, Clone MI15) to stratify them into lymphoid, myeloid and pauci-immune pathotypes according to previously published histological features *(30, 78)*. Fibrosis was assessed by Elastica van Gieson staining. Acute inflammation was assessed by staining for neutrophils/CD15 *(29)*. Crystals were sought on the biopsy to rule out a microcrystalline arthritis.

**Processing of synovial tissue biopsies**

scRNA-sequencing was performed in seven SSc biopsies, six RA biopsies, and three NIC samples as described before *(79)*. Additional three NIC samples were publicly available on ArrayExpress (E-MTAB-14339). Synovial biopsies were washed with phosphate-buffered saline, mechanically minced and enzymatically digested using Liberase TL (100 µg/mL; Roche) and DNase I (100 µg/mL; Roche) in RPMI 1640 cell culture medium (Thermo Fisher) for 30 min at 37°C. After stopping the digestion process with fetal calf serum (FCS), erythrocytes were lysed with Red Blood Cell Lysis solution (Miltenyi Biotec). Cells were washed and counted on an automated cell counter (LUNA Logos Biosystems or Invitrogen Countess® III FL).

**Single-cell RNA sequencing (scRNA-seq)**

A total 6000 unsorted synovial cells per patient were targetetd and prepared for scRNA-seq analysis using the Chromium Single Cell 3' GEM, Library & Gel Bead Kit v3, the Chromium Chip B Single Cell Kit (10X Genomics) and the Chromium controller (all 10X Genomics). Libraries were sequenced at the Functional Genomics Centre Zurich (FGCZ) on the Illumina NovaSeq instrument, demultiplexed using CellRanger (v7.2.0) from 10X Genomics and mapped to the reference genome GRCh38.p13.

**Analysis of scRNA-seq data**

Analysis of the scRNA-seq data was mainly performed in R (v4.3.1), using Seurat (v4.3.0.1) *(80)*. Raw samples were corrected for ambient RNA using SoupX (v1.6.2) *(81)*. All samples were quality-controlled in the same way: (1) discarding the top 1% of features and counts, (2) cells with less than 200 features, (3) the top 5% or if exceeding more than 20% mitochondrial gene content, (4) doublets estimated by scDblFinder (v1.15.2) *(82)*.

The log-normalised expression matrices across all biopsies were merged and scaled, regressing out the number of features, number of counts, mitochondrial content and cell cycle effects as implemented by Seurat. PCA was performed on the top 3000 most variable genes shared across the samples, picking the optimal number of dimensions as determined by the elbow method. Louvain clustering on the shared nearest-neighbor (SNN) graph was performed on the batch-corrected dimensions obtained by Harmony (v0.0.1) *(83)*. Using a resolution of 0.05 with stable basic clusters as defined by clustree (v0.5.0) *(84)* we performed manual cell type annotation using established marker genes. The four largest cell populations (fibroblasts, myeloid cells, endothelial cells, T cells) were reclustered for annotation of subpopulations. Each cell type was iteratively integrated separately with STACAS (v2.2.2) *(85)*, ensuring to exclude low-quality cells (defined by expression of non-cell type markers and clustering by technical batch effect, apoptotic markers). Final subpopulations were annotated based on unsupervised clusters of the SNN graph, taking into account established markers from previous studies *(16, 17, 31)*. Proportion differences were statistically analysed by a pairwise Wilcoxon rank sum test as implemented by pochi (v0.1.0) *(86)* and a permutation test using scPropotionTest (v0.0.0.9000) *(87)*.

Pairwise differential expression analysis (DEA) between diseases was performed using Seurat's implementation of MAST *(88)* on genes expressed in >20% of cells. Genes were defined as differentially expressed at a Benjamini-Hochberg (BH) corrected p-value of < 0.01 and absolute log2FC > 0.25. Over-representation analysis (ORA) was performed using clusterProfiler (v4.10.1) *(89)* using gene sets from different collections (MSigDB hallmark *(90)*, Gene ontology, Reactome, KEGG, WikiPathways) and manually curated lists of genes from literature as indicated in the main text. Gene set scores were calculated using UCell (v2.14.0) *(91)*. Differentially expressed genes in SSc SF (vs. RA) contributing to the Hallmark ORA were subjected to STRING *(92)* network enrichment analysis (Markov Cluster Algorithm (MCL), inflation parameter = 2) to isolate functional modules from overlapping gene sets. Activity inference was conducted with decoupleR (v2.8.0) *(93)* using a multivariate linear model for pathways (PROGENy) *(94)* and a univariate linear model for transcription factor activity estimation (CollecTRI) *(95)*. Cell-cell interaction analysis was performed with CellChat (v 2.1.2) following standard parameters *(96)*.

Plots were generated using scCustomize (v1.1.3) *(97)*, SCpubr (v2.0.2) *(98)*, pathview (v1.42.0) and custom ggplot2 (v3.5.1) scripts.

**Spatial transcriptomics**

Synovial tissue from three SSc patients was processed into 5 μm formalin-fixed paraffin-embedded (FFPE) sections for single-cell spatial transcriptomics using the 10x Genomics Xenium platform. Gene expression was profiled using a 477-gene panel comprising the Xenium Multi-tissue and Cancer Panel (377 genes) supplemented with a 100-gene custom add-on (Table S5) targeting cell subpopulations and IFN response genes selected from the previous scRNA-seq analysis. Data were acquired using Xenium instrument software v3.4.1.0 and processed with Xenium analysis pipeline v3.3.0.1.

Transcript counts were imported into R (v4.5.2) using Seurat (v5.3.1) *(99)*. Synovial tissue regions of interest were manually defined in XeniumExplorer (v4.1.1) and exported per tissue piece. Cells were retained if they exceeded minimum thresholds for transcript count (> 10) and detected features (> 5), and cells in the top 1% of transcript counts were excluded as potential doublets or segmentation artefacts. Gene expression was log-normalised (scale factor 10,000).

For dimensionality reduction, all 477 panel genes were used after scaling with regression of total transcript and feature counts. PCA was computed on 30 components and a shared nearest neighbour graph was constructed for Louvain clustering across a resolution sweep (0.5–1.2), with the optimal resolution selected per sample based on cluster stability (clustree v0.5.1). Louvain-derived clusters were then manually annotated per patient based on marker gene expression and visually confirmed in the spatial coordinates. To resolve fibroblast and myeloid subpopulations, RCTD (spacexr v1.2.0) *(100)* deconvolution was applied separately to these lineages, using the SSc synovial scRNA-seq dataset described above as reference (one of seven scRNA-seq patients was also included in the spatial cohort). RCTD-derived subtype labels were integrated with the manual annotation into a unified per-cell label. As the granularity of resolvable populations varied across patients, labels were subsequently harmonized across the three samples. For main figures, sublining SF and dendritic cell popualtions were collapsed into broader categories. For multi-sample visualisation, objects were merged and a Harmony-corrected embedding was computed for UMAP visualization only; all biological analyses used uncorrected expression data.

IFN response gene sets were obtained from the MSigDB Hallmark collection *(90)* and intersected with the Xenium panel. Per-cell IFN programme scores were computed using UCell (v2.14.0) *(91)*, which applies a rank-based scoring approach ensuring cross-sample comparability without additional normalisation. To characterise transcriptionally distinct IFN programmes, mean scaled expression of panel-intersecting Hallmark IFN genes was computed per cell type, Z-scored, and subjected to hierarchical clustering (ward.D2), identifying a stromal and a myeloid IFN programme.

Spatial autocorrelation of program-specific UCell scores was assessed per tissue piece using global Moran's I, computed on a k-nearest neighbour spatial weight matrix (k = 15) implemented in spdep (v4.1-1). Local spatial clustering was characterised using Local Moran's I (LISA), classifying cells into High-High (HH; score above global mean, positive local Moran's I, $p < 0.05$), Low-Low (LL; score below global mean, positive local Moran's I, $p < 0.05$), spatial outlier ($p < 0.05$, remaining), or non-significant quadrants. All spatial statistics were computed per tissue piece to avoid coordinate system confounding. Cell type enrichment in HH versus LL niches was quantified as log2 odds ratio per patient, with a pseudocount of 0.5 applied to prevent undefined values when a cell type was absent from a niche.

**Spatial proteomics using sequential immunofluorescence (seqIF)**

Protein-level validation was performed on 5 µm FFPE sections from four SSc patients, including consecutive sections from the three patients profiled by spatial transcriptomics, using the Lunaphore COMET sequential IF platform with a 33-plex panel optimised across eight development runs, targeting synovial cell populations and IFN response markers (antibody details in Table S6).

Sections were manually dewaxed by sequential immersion in xylene (2 × 5 min) and 100% ethanol (2 × 2 min), followed by drying at 60°C for 5 min. Antigen retrieval was performed in EDTA buffer (pH 9) at 99°C for 60 min using a PT-Module (Epredia). Non-specific binding was blocked with 5% horse serum in multi-staining buffer (MSB, Lunaphore) for 30 min at room temperature. Cyclic immunofluorescence was performed across 18 antibody cycles on the COMET platform. Autofluorescence images were acquired before the first cycle and after every one to four cycles for background subtraction. DAPI was acquired each cycle for image registration. Images were analysed using Lunaphore HORIZON software (v2.5). Nuclear localisation of pSTAT1, pSTAT2, and IRF1 was assessed as a readout of IFN transcription factor activity at the protein level. MX1 expression was evaluated as a marker of type I IFN pathway activation. All data were acquired at the UZH Center for Microscopy and Image Analysis (ZMB).

Cell-type annotation was performed using CellTypist (v1.7.1) *(101)*, applying tissue-appropriate reference models to each tissue object. Specifically, the Adult_Human_Skin model to skin dataset, and the Immune_All_High model to the PBMCs dataset.

Type I and II IFN pathway activation scores were calculated from MSigDB Hallmark gene sets using pyUCell (v0.6.0) *(102)* by default parameters, the Z-score was then computed per cell type per sample relative to controls. The ability of IFN Z-scores to discriminate SSc from control samples was validated by hierarchical clustering (Euclidean distance, complete linkage).

**Bulk RNA-seq of synovial fibroblasts**

SF were isolated from synovial tissues obtained during synovectomy or joint replacement surgery from five RA patients fulfilling the ACR/EULAR classification criteria *(69)*. SF were isolated and passaged in RPMI1640 supplemented with 10% FCS and 1% penicillin/streptomycin (all Gibco) until passage 3, then used for experiments up to passage 9 $1\times10^5$ SF were seeded in 1 mL complete medium in 24-well plates. After 24 hours, SFs were treated with 10 ng/mL IFN-β or IFN-γ (both R&D). After 24 hours of treatment, RNA was isolated using the RNeasy Mini Kit (Qiagen) according to the manufacturer's protocol. RNA-Seq was performed using QuantSeq 3' mRNA-Seq Library Prep Kit FWD for Illumina (Lexogen) according to the manufacturer's protocol. The libraries were sequenced as 50 bp SE on an Illumina HiSeq instrument.

TNF-stimulated SF cultures were previously described in *(103, 104)*. Raw read quality was evaluated using FastQC. Reads were aligned to the hg38 assembly and quantified using STAR (v2.5.4b). ComBat_seq (v3.42.0) was used to adjust for batch effects and DESeq2 (v1.34.0) was used to normalize gene counts and perform differential expression analysis.

Additionally, publicly available bulk RNA-seq data from IFN-α, IFN-γ and TNF-stimulated synovial fibroblasts were obtained from Tsuchiya et al. *(105)* and used as independent validation of cytokine-specific transcriptional stimulation signatures.

**CD55 immunohistochemistry and scoring**

FFPE synovial tissue sections from our cohort of patients with SSc or RA were analysed for CD55 expression in the synovial lining. CD55 was detected using a monoclonal anti-CD55 antibody (Abcam, ab133684), with staining performed on a Leica BondMAX automated immunostainer. Antigen retrieval followed the Leica H1 antigen retrieval protocol (20 min, 95°C). Antibody titration was performed on synovial tissues from RA (n = 2) and osteoarthritis (OA, n = 2) patients using serial dilutions (1:4,000 to 1:12,000) to identify a working concentration that enabled discrimination between weak and strong staining. A final dilution of 1:8,000 was selected for subsequent analysis.

The cohort included FFPE blocks from a subset of the study cohort described above (11 SSc and 6 RA patients). Three SSc samples were excluded due to insufficient tissue or unclear identification of synovial lining, resulting in a final dataset of 9 SSc and 7 RA samples. Only regions with a clearly identifiable synovial lining layer were evaluated. Staining intensity was scored semi-quantitatively by two independent, blinded observers (CG; CCC) using a 4-point scale (0 to 3, in 0.5-point increments). Inter-rater reliability was high, with an intraclass correlation coefficient (ICC) of 0.931 (95% CI: 0.815-0.974).

***Ex vivo* SF culture and stimulation**

Synovial tissue samples were collected from NIC or OA patients undergoing joint replacement surgery at Schulthess Clinic Zurich, Switzerland or via ultrasound-guided biopsy at University Hospital Zurich and Inselspital, University Hospital Bern, Switzerland.

Fresh synovial tissue was processed as described above. The resulting cell suspension was resuspended in complete DMEM (DMEM supplemented with 10% FCS, 2 mM L-glutamine, 10 mM HEPES, 0.2% amphotericin B and 50 U/mL penicillin/streptomycin). Cells were maintained at 37°C in an incubator containing 5% $CO_2$. SF from passages 4 to 7 were used for subsequent experiments.

2D-cultured SFs were seeded and cultured 24 h prior to stimulation with human recombinant IFN-α2a (10 ng/mL; ImmunoTools), human recombinant IFN-β1a (10 ng/mL; ImmunoTools), human recombinant IFN-γ, (10 ng/mL; ImmunoTools), or  human recombinant TNF (10 ng/mL; R&D Systems) and cultured for 8, 24 or 48 hours.

3D synovial micromasses were created by resuspending SFs from OA or NIC patients in a single 35-µL droplet of Matrigel (Matrigel® Growth Factor Reduced Basement Membrane Matrix, Phenol Red-Free, Corning, Cat. No. 356231) at a concentration of $2x10^5$ cells per organoid. Co-cultures resuspended in Matrigel were seeded in a single well of polyHEMA (Sigma-Aldrich, Cat. No. 192066-10G)-coated plates and were left for 1 hour in the incubator at 37°C to polymerize before adding complete DMEM medium. 3D micromasses were stimulated three times per week with IFN-α2a, IFN-γ or TNF for 14 days at the same concentration as described for 2D SF cultures.

**Quantitative real-time polymerase chain reaction (qPCR)**

RNA was extracted from the stimulated SFs using the RNeasy plus Mini Kit (Qiagen) following the manufacturer’s guidelines, with subsequent cDNA generation using the cDNA Synthesis Kit

(Thermo Fisher Scientific). FastStart Universal SYBR Green MM (Roche) was used for qPCR. The list of the primer pairs used in this study is provided below:

| **Gene** | **Forward Primer (5' → 3')** | **Reverse Primer (5' → 3')** |
|---|---|---|
| GAPDH | GGG AAG CTT GTC ATC AAT GGA | TCT CGC TCC TGG AAG ATG GT |
| CD55 | ACC AAA TGC TCA AGC AAC ACG | GCT TGG TTG TCC TGG AAA CAG |
| CFB | AAG GCA GAG TGC AGA GCA AT | GTT CTC GAA GTC GTG TGG TCT |
| CFD | GGA CAG CTG CAA GGG TGA CTC | TGA CCA CGC CCT CGA GCA |
| CFH | AAG CGC AGA CCA CAG TTA CAT | CTT GAT TTG GAA CAT GTT TTG ACA C |
| CFI | AAC TGT GTT GTA AAG CAT GCC A | CCC GAT TTG CAA TGG AAG CC |
| CLU | GCC AAG AAG AAG AAA GAG GAT GC | GTC TCT GAT TCC CTG GTC TCA TT |
| C3 | ACA GCT GAA GGA AAA GGC CA | GGT ACA TTG TCA CCA CCG ACA |
| SERPING1 | CCA GAA GTT TGG AGT CCG CT | GCA TTT GGA TTT GAG GAG GCT |
| COMP | GCA GAC AAT GAA CAG CGA CC | TGA AGG CAG TGT AAC CCA CA |

Data were analysed using the comparative CT method and expressed as $2^{-\Delta\Delta CT}$ with GAPDH as internal standard for sample normalization. Fold change values were log2-transformed prior to statistical analysis. Outliers were identified and removed using the ROUT method (Q = 1%). To assess whether each stimulation condition resulted in significant changes in gene expression compared to unstimulated controls, a two-tailed one-sample t-test against a theoretical mean of zero was performed for each condition and gene separately. Statistical analyses were performed using GraphPad Prism (v11.0.1). A p-value < 0.05 was considered statistically significant.

### ELISA

Cell culture supernatants were tested using the Complement Factor C3a (ab279352), Complement Factor B (Ba fragment, ab315062), and Complement Factor H (ab252359) ELISA kits (all

Abcam). SF micromass supernatants were tested with the human Total MMP-1 DuoSet ELISA kit (R&D Systems, DY901B-05) and Total MMP-3 DuoSet ELISA kit (R&D Systems, DY513-05). ELISA data were analyzed using a one-way ANOVA with Dunnett's test versus non-stimulated control. Statistical analyses were performed as described above for qPCR.

**Statistical assessment of IFN-α2a serum levels as predictor for SSc synovitis**

The cohort of 132 SSc patients for this analysis was derived from Gerges et al. *(54)*, excluding patients with overlapping diseases (Table S3). Joint involvement was assessed clinically and categorised as either swollen joints (synovitis), tender joints, both swollen and tender (synovitis), or none. Serum IFN-α2a concentrations (pg/mL) were $\log_{10}$-transformed to correct for right-skewed distribution. We first applied a binary logistic regression to test whether $\log_{10}$ IFN-α2a levels were associated with the presence of swollen joints (binary outcome). The area under the receiver operating characteristic curve (AUC) was calculated from the binary logistic regression model using the pROC package (v1.18.5). To explore differential associations across types of joint involvement, a multinomial logistic regression model was fitted using joint phenotype as a three-level categorical outcome variable ("none", "swollen", "tender only"), with "none" as the reference category. Odds ratios and 95% confidence intervals were calculated, and statistical significance was defined as $p < 0.05$. All analyses were performed in R (v4.3.1) using the stats and nnet packages (v7.3-19).

**Lentiviral ISRE-reporter SF cell line generation and serum stimulation assay**

The pGreenFire1-ISRE lentiviral reporter plasmid (System Biosciences), encoding an ISRE-driven destabilized copGFP and luciferase reporter with EF1α-driven puromycin resistance (pGreenFire1-ISRE EF1α-Puro, TR016PA-P), was transformed into NEB Stable competent *E. coli*

(C3040I) according to the manufacturer's protocol. Plasmid DNA was purified using the ZymoPURE II Plasmid Midiprep Kit (Zymo research, D4201-A).

Lentiviral particles were produced in HEK293T cells using a second-generation packaging system. HEK293T cells were co-transfected with pGreenFire1-ISRE, psPAX2 and pMD2.G at a 2:1:1 molar ratio using Lipofectamine 3000 (Thermo Fisher Scientific) according to the manufacturer's instructions. Viral supernatants were collected at 24 and 48 hours after transfection and stored at –80°C to be used for downstream transduction experiments.

Functional viral titres were determined by transducing HEK293T cells with serial dilutions of viral supernatant in the presence of polybrene (6 µg/mL, TR-1003-G, Merck). Cells were incubated with virus-containing medium for 8 hours at 37°C, after which the medium was replaced with DMEM. Reporter activity was validated by stimulating transduced cells with IFN-α2a (10 ng/mL, ImmunoTools) and measuring destabilized copGFP expression at 8 and 16 hours by flow cytometry on a BD LSRFortessa instrument (Cytometry Facility, University of Zurich). Data were analyzed using FlowJo (v10.10.0).

Primary OA synovial fibroblasts from two donors were transduced with the same lentiviral batch at a multiplicity of infection (MOI) of 3. Virus-containing medium supplemented with polybrene (6 µg/mL, Sigma-Aldrich) was added to the cells, followed immediately by spinoculation at 1500 g for 90 min at 32°C. The virus-containing medium was then removed without further incubation or washing, fresh DMEM was added, and cells were cultured at 37°C.

Forty-eight hours after transduction, cells were selected with puromycin (3 µg/mL, TargetMol) for 3 days to establish two stable pGreenFire1-ISRE reporter synovial fibroblast lines.

To optimise assay conditions, reporter cells were stimulated with human recombinant IFN-α2a (ImmunoTools) under different concentrations (10 ng/mL, 10 pg/mL, 10 fg/mL) and time conditions (8 h, 16 h), and reporter activity was assessed 6 and 24 h after stimulation. Luciferase activity and cell viability were measured using the ONE-Glo™ + Tox Luciferase Reporter and Cell Viability Assay Kit (E7110, Promega). Fluorescence and luminescence signals were recorded using an Agilent BioTek Synergy H1 multimode reader. Luciferase values were normalized to the corresponding fluorescence-based viability signal to obtain the final reporter output. As the 6-hour stimulation timepoint produced overall stronger reporter induction than the 24-hour timepoint, subsequent experiments were performed using the 6 h stimulation using 10% SSc patient serum (n = 38) or healthy serum (n = 4, sex- and age-matched) diluted in DMEM (Table S4). Luciferase activity was normalised to unstimulated controls to obtain fold change values. Differences between groups were assessed using a Wilcoxon rank-sum test with BH correction.

**Comparative analysis of SLE arthritis and Anifrolumab response signatures**

The 29-gene signature specific to SLE arthritis relative to OA and RA was obtained from *(59)*. Anifrolumab response genes were obtained from supplememtary data of *(58)*, including 526 genes downregulated in skin biopsies at day 28 and mapped to gene symbols using biomaRt (v2.85.2). For comparison with these microarray-derived signatures, SSc scRNA-seq data from each biopsy was collapsed into pseudobulk using Seurat's *AggregateExpression()* function. We then performed differential expression analysis with DESeq2 (v1.50.2) and GSEA using those signatures on pseudobulk-aggregated data from SSc synovium using clusterProfiler (v4.18.4) *(89)*.

**Publicly available SSc skin and PBMC scRNA-seq data processing**

To compare SSc synovium and skin transcriptomics, publicly available scRNA-seq datasets on SSc skin were retrieved from GEO and ArrayExpress (GSE138669 and GSE209635) *(106, 107)*. Similarly the peripheral blood mononuclear cell (PBMC) public data were retrieved from one dataset (E-GEAD-872) *(48)*. For both the skin and PBMC datasets, each dataset was preprocessed independently using Scanpy (v1.11) *(108)*. Quality control was performed by applying dynamic thresholds on the following per-cell metrics: mitochondrial RNA fraction ($95^{th}$ percentile or < 20%), total UMI count (> 200), number of detected genes or UMI ($99^{th}$ percentile). Genes with less than 10 reads were excluded from the analysis. Doublets were identified and removed using Scrublet *(109)*. Gene expression counts were normalized to 10,000 counts per cell followed by log1p transformation.

Independently for the skin-synovium and PBMC objects highly variable genes were selected, and the number of genes and ribosomal and mitochondrial RNA content regressed out. Principal component analysis (PCA) was performed on the scaled data, followed by Harmony integration (harmonypy, v0.0.10) with default parameters *(110)*. The neighbourhood graph was constructed using the Harmony-corrected factors with n_neighbors = 15 and n_pcs = 50, followed by UMAP dimensionality reduction.

**Figures**

Figures created with BioRender were licensed as follows:

Fig. 1A (https://BioRender.com/av1evel)

Fig. 7 (https://BioRender.com/2u3k7xy)

Fig. S4C (https://BioRender.com/oh26355)

**Use of artificial intelligence tools**

Large language models (ChatGPT, OpenAI; and Claude, Anthropic) were used to assist with manuscript writing and editing, coding, and scientific discussion. All AI-assisted content was critically reviewed, verified, and edited by the authors, who take full responsibility for the accuracy and integrity of the work.

**Study approval**

The collection and experimental usage of the human samples was approved by the ethical commission of the Kanton Zurich (approval numbers: BASEC 2019-00674, 2021-00092, 2021-01818). Written informed consent was obtained from all patients. All experiments were performed in accordance with the institutional guidelines.

**Patient research partner engagement**

We engaged two patient research partners, A. Eisenring and J. Messmer, throughout the study, ensuring that our research questions and outcomes are directly aligned with the needs and expectations of those affected by SSc.

**Data availability**

RNA-seq, scRNA-seq, and spatial transcriptomics data generated in this study will be deposited in the NCBI Gene Expression Omnibus (GEO) or ArrayExpress upon manuscript acceptance. Accession numbers will be provided upon publication. The following publicly available datasets were used in this study: NIC synovial scRNA-seq data (E-MTAB-14339 and E-MTAB-16064, ArrayExpress); SSc PBMC scRNA-seq data (E-GEAD-872, ArrayExpress); SSc skin scRNA-seq data (GSE138669 and GSE209635, GEO).

Any additional information required to reanalyze the data reported in this paper is available from the lead contact upon request.

### Code availability

The code used for the scRNA-seq and spatial transcriptomics analysis will be made available on GitHub upon publication.

### SUPPLEMENTARY FIGURES

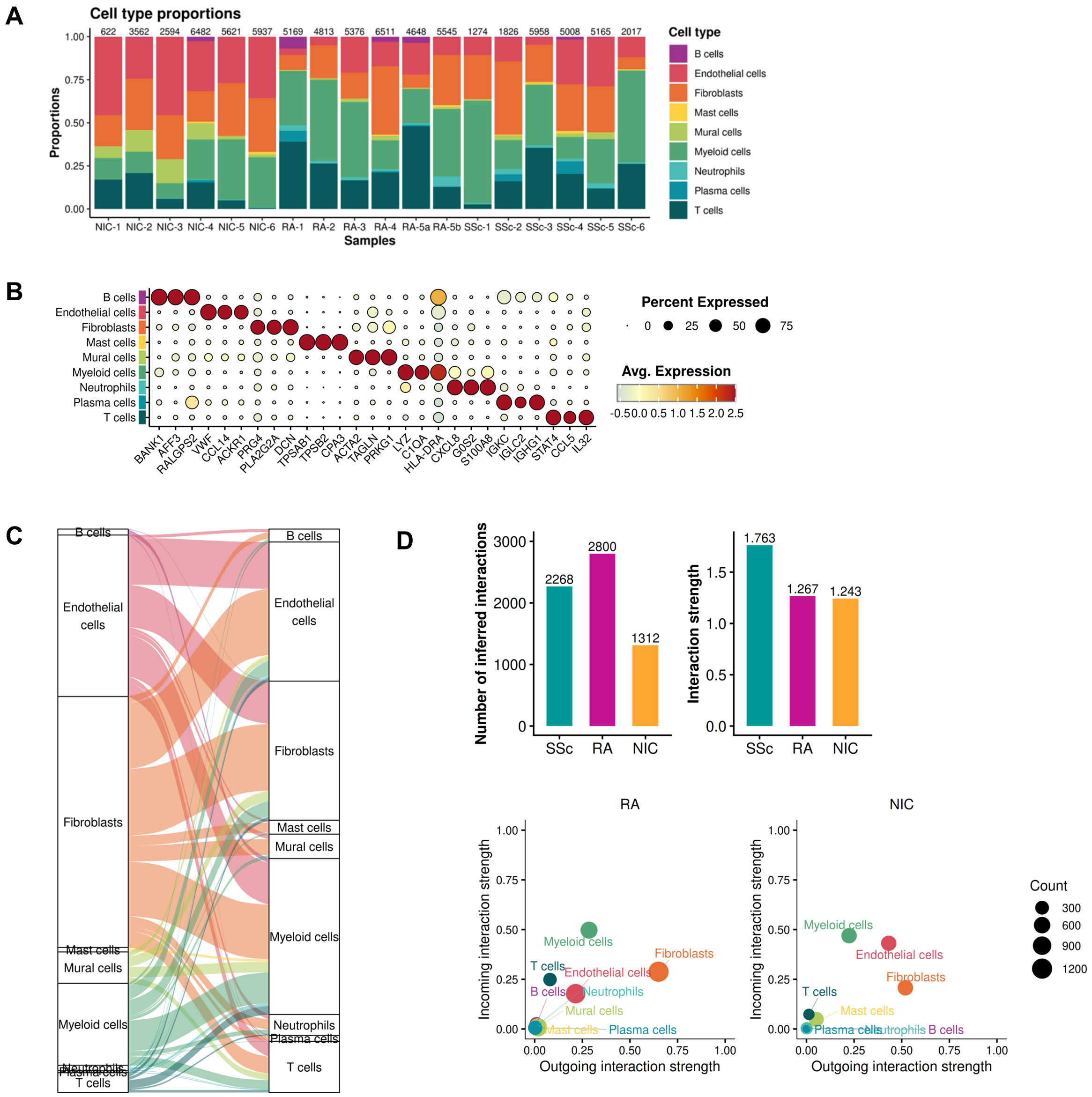


**Fig. S1. Synovial cell type composition and intercellular communication across SSc, RA, and NIC.** (**A**) Cell type proportions per biopsy; numbers above bars indicate cells per sample. (**B**) Top three marker genes per cell type; dot size indicates the percentage of expressing cells and color the average expression. (**C**) Cell-cell interactions between cell types in SSc (CellChat). (**D**) Number of inferred interactions and overall interaction strength in SSc, RA, and NIC (CellChat). (**E**) Incoming and outgoing interaction strength per cell type in RA and NIC (CellChat); compare Fig. 1H (SSc).

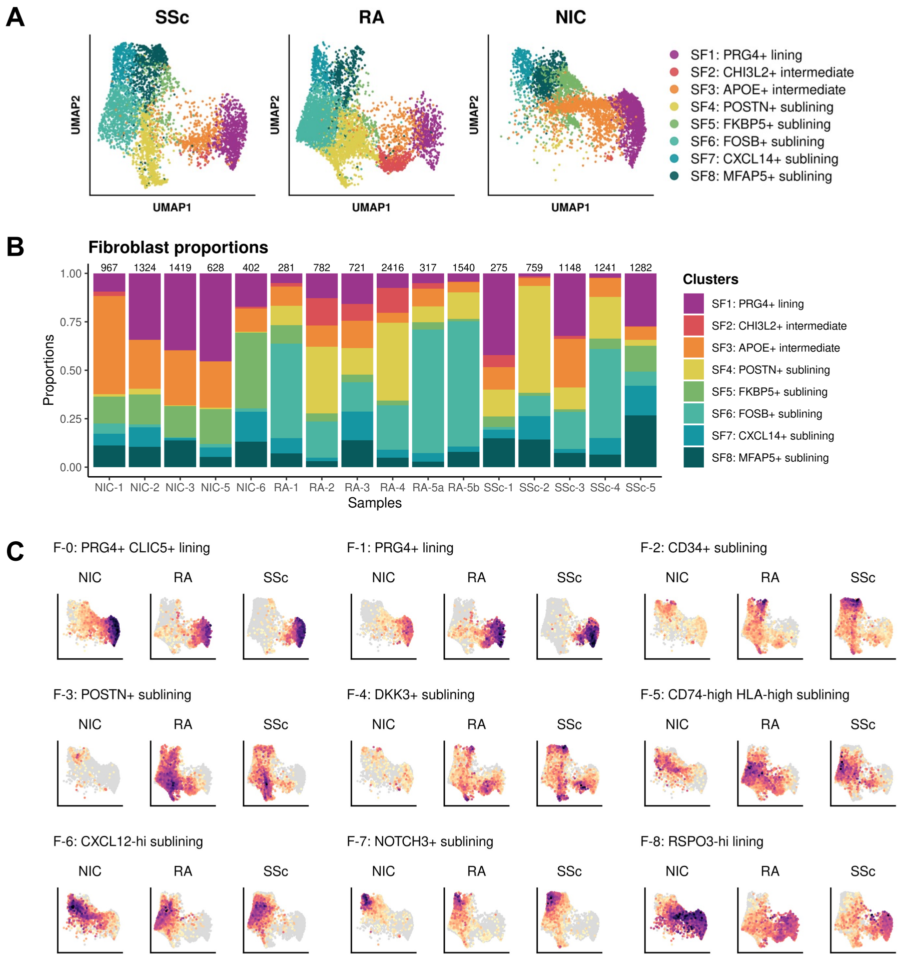


**Fig. S2. Synovial fibroblast subpopulations across SSc, RA, and NIC.** (**A**) UMAP of SF populations (SF1–SF8) split by disease group. (**B**) Proportions of SF populations per biopsy; numbers above bars indicate cells per sample. (**C**) Seurat module scores of top ten fibroblast population markers from Zhang et al. 2023 (F-0 to F-8) across the SF UMAP.

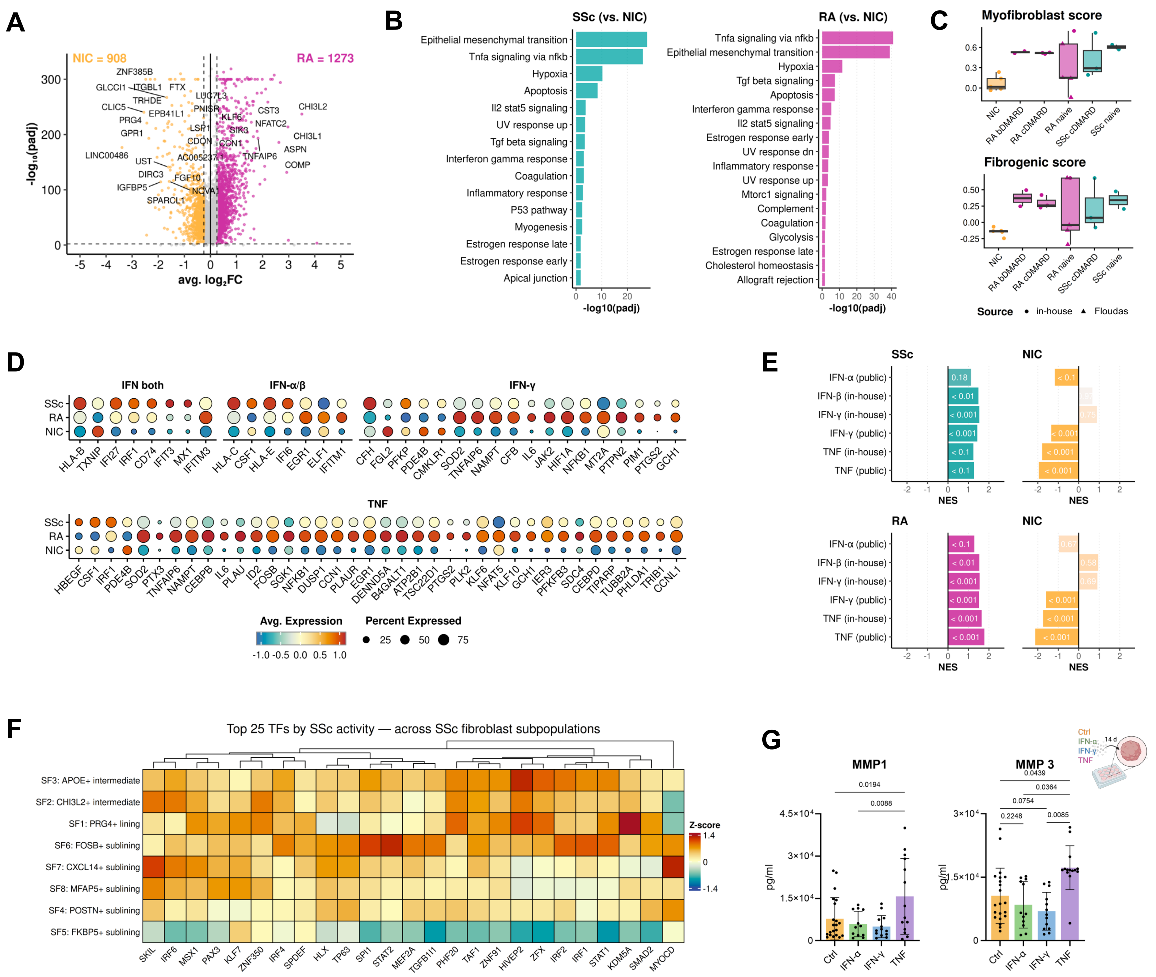

**Fig. S3. SSc vs. RA SF transcriptional programs.** (**A**) Volcano plot of differentially expressed genes (DEGs) in RA vs. NIC SF; numbers indicate up-regulated genes per group. (**B**) Pathway over-representation analysis of SSc and RA DEGs versus NIC (MSigDB Hallmark). (**C**) Synovial myofibroblast and fibrogenic scores including additional treatment-naïve RA (n = 4, Floudas); symbols denote source dataset. (**D**) IFN (both, IFN-α/β, IFN-γ) and TNF response DEGs between SSc and RA; dot size indicates the percentage of expressing cells, color the average expression. (**E**) GSEA of SSc, RA, and NIC DEGs against ranked expression of in vitro–stimulated SF, extending Fig. 3I to the NIC comparison. (**F**) Top 25 transcription factors by SSc activity across

SF subpopulations (decoupleR, CollecTRI). (**G**) MMP1 and MMP3 ELISA of SF micromass supernatants after 14 days of stimulation (one-way ANOVA with Tukey post-hoc test).

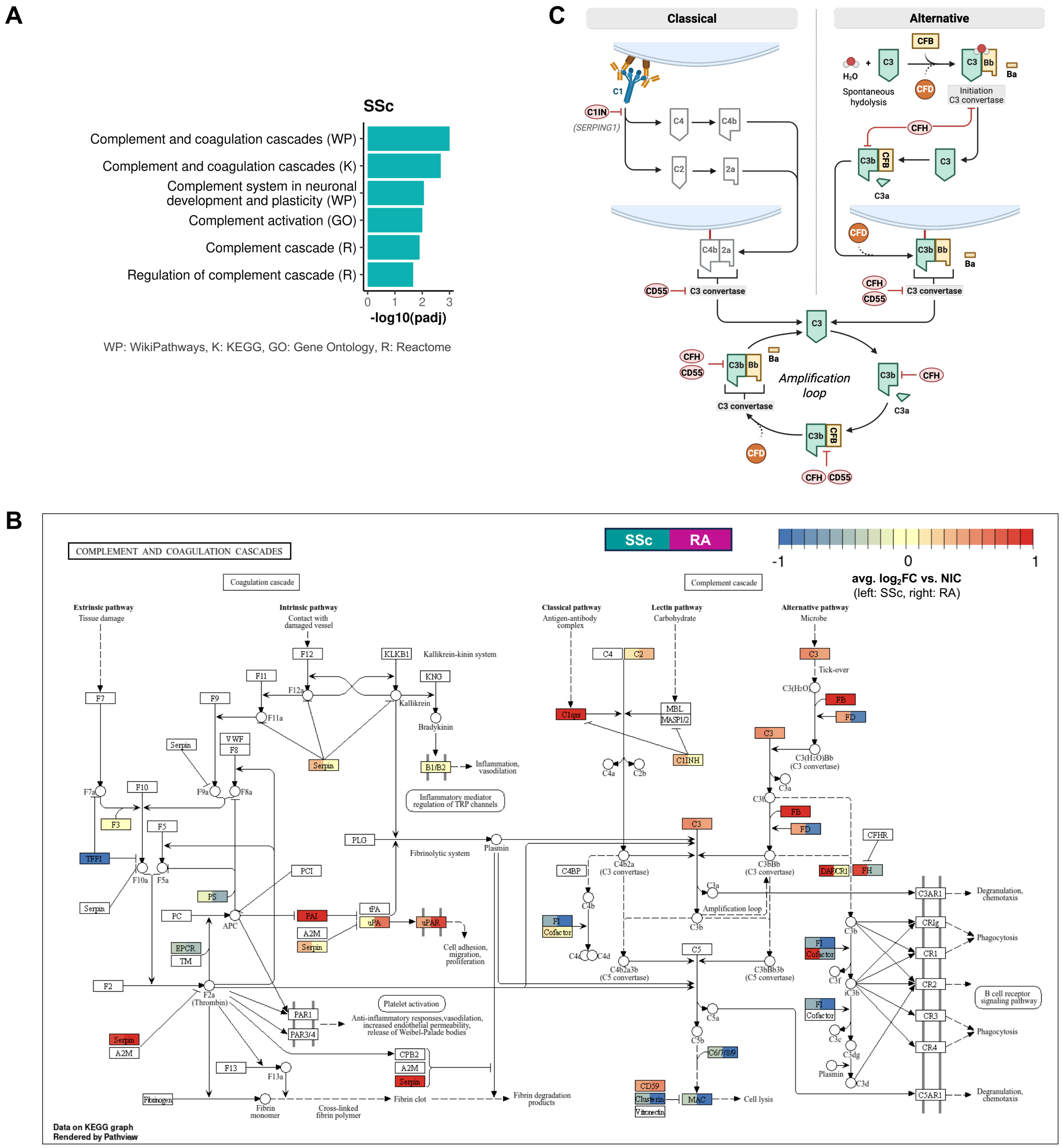


**Fig. S4. Complement pathway dysregulation in SSc synovial fibroblasts.** (**A**) Pathway over-representation analysis of SSc vs. RA DEGs enriched in complement and coagulation gene sets across multiple. (**B**) KEGG complement and coagulation cascade map; node color indicates average log2FC versus NIC (left half SSc, right half RA), white indicates no detected expression.

(**C**) Summary schematic of complement regulation in SSc; genes with increased expression in SSc ($log_2FC > 0.25$ versus NIC) in color, C2 and C4 in white. Created in BioRender.

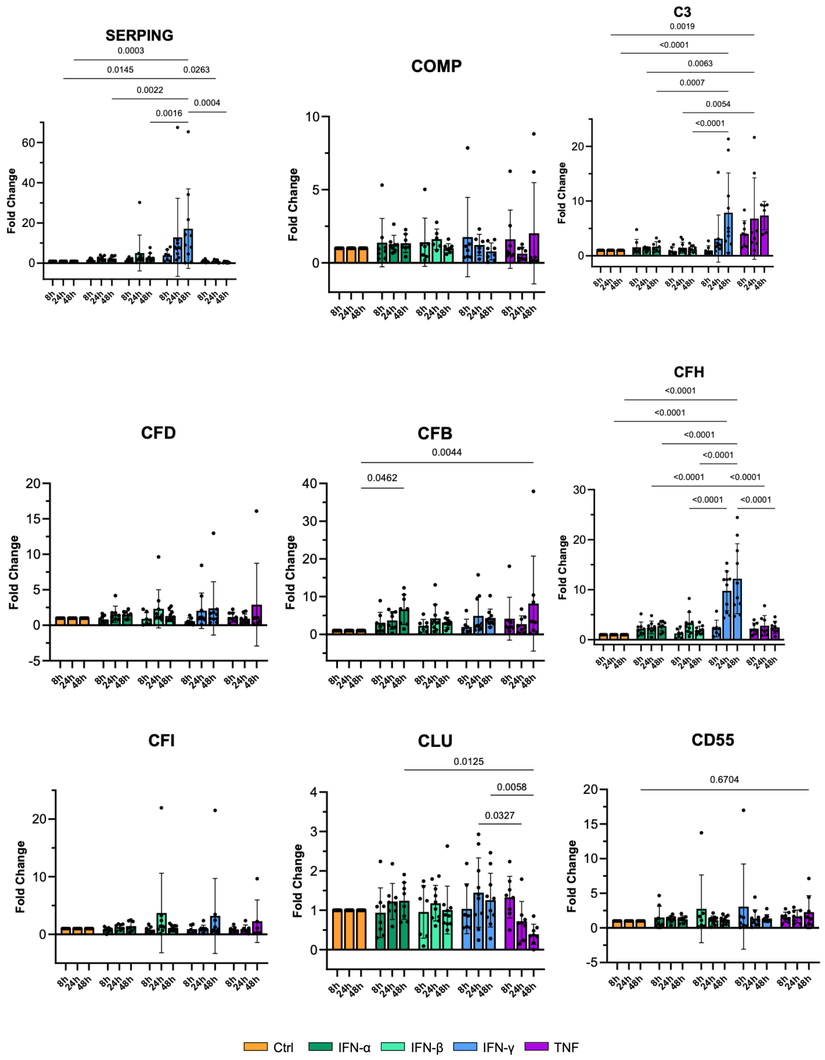


**Fig. S5. Time course of complement gene expression in IFN- and TNF-stimulated synovial fibroblasts.** Fold change in complement gene expression (RT-qPCR) in SF after 8, 24, and 48 h of stimulation with IFN-α, IFN-β, IFN-γ, or TNF (10 ng/ml each, 24 h); compare Fig. 4G. One-way ANOVA with Tukey post-hoc test; n = 8-10.

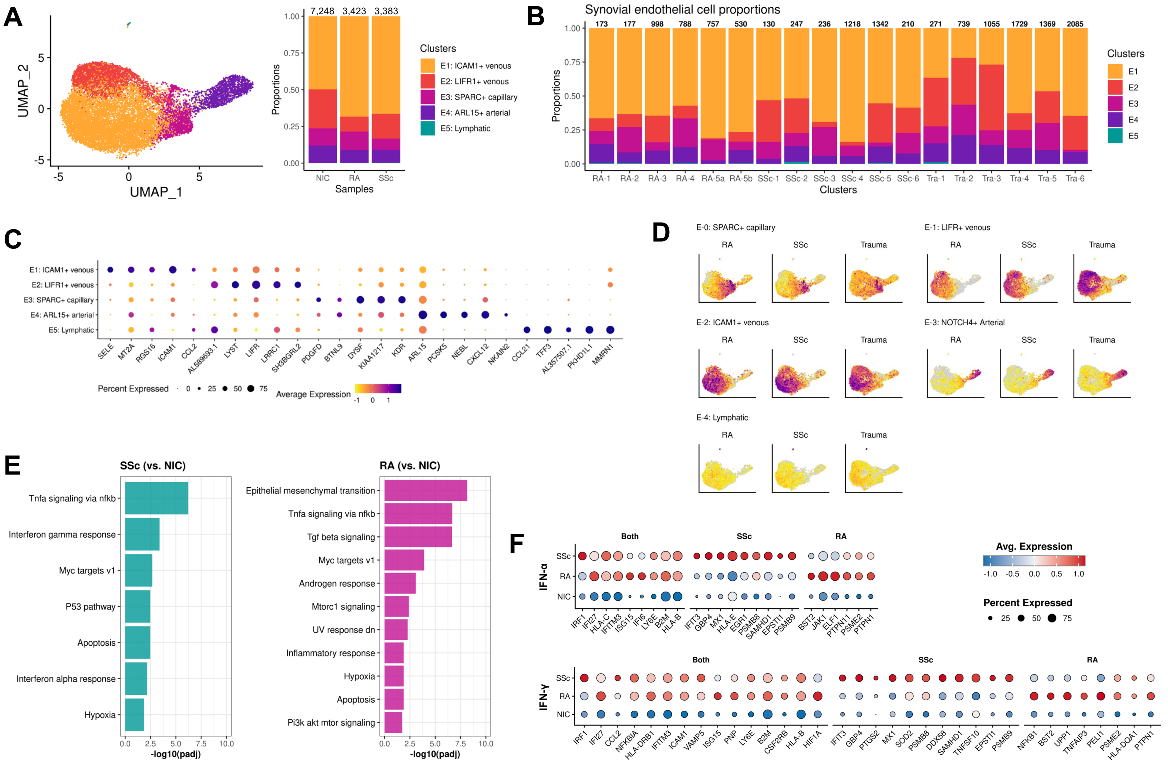


**Fig. S6. Characterization of endothelial cell (EC) populations.** (**A**) UMAP of re-integrated EC populations split by disease group. (**B**) Proportions of EC populations per biopsy; numbers above bars indicate cells per sample. (**C**) Top five marker genes per EC population. (**D**) Module scores of published EC subpopulation signatures (Zhang et al. 2023, E-0 to E-4) projected onto the EC UMAP, split by disease group (Trauma = NIC). (**E**) Pathway over-representation analysis of SSc and RA DEGs versus NIC (MSigDB Hallmark). (**F**) Differentially expressed Hallmark IFN-α and IFN-γ response genes of SSc vs. NIC (SSc), RA vs. NIC (RA) or both SSc/RA vs. NIC (Both).

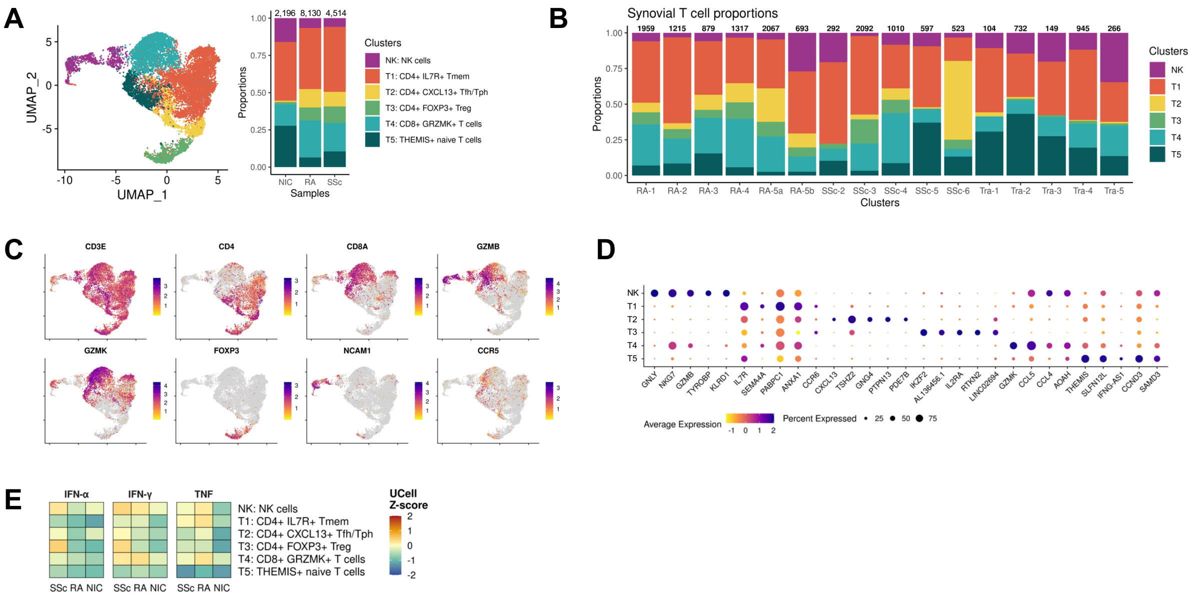


**Fig. S7. Characterization of T cell populations.** (**A**) UMAP of re-integrated T cell populations split by disease group. (**B**) Proportions of T cell populations per biopsy; numbers above bars indicate cells per sample. (**C**) Expression of canonical T cell marker genes across the T cell UMAP. (**D**) Top five marker genes per T cell subpopulation. (**E**) Hallmark IFN-α, IFN-γ, and TNF response gene set scores across endothelial cell populations (UCell).

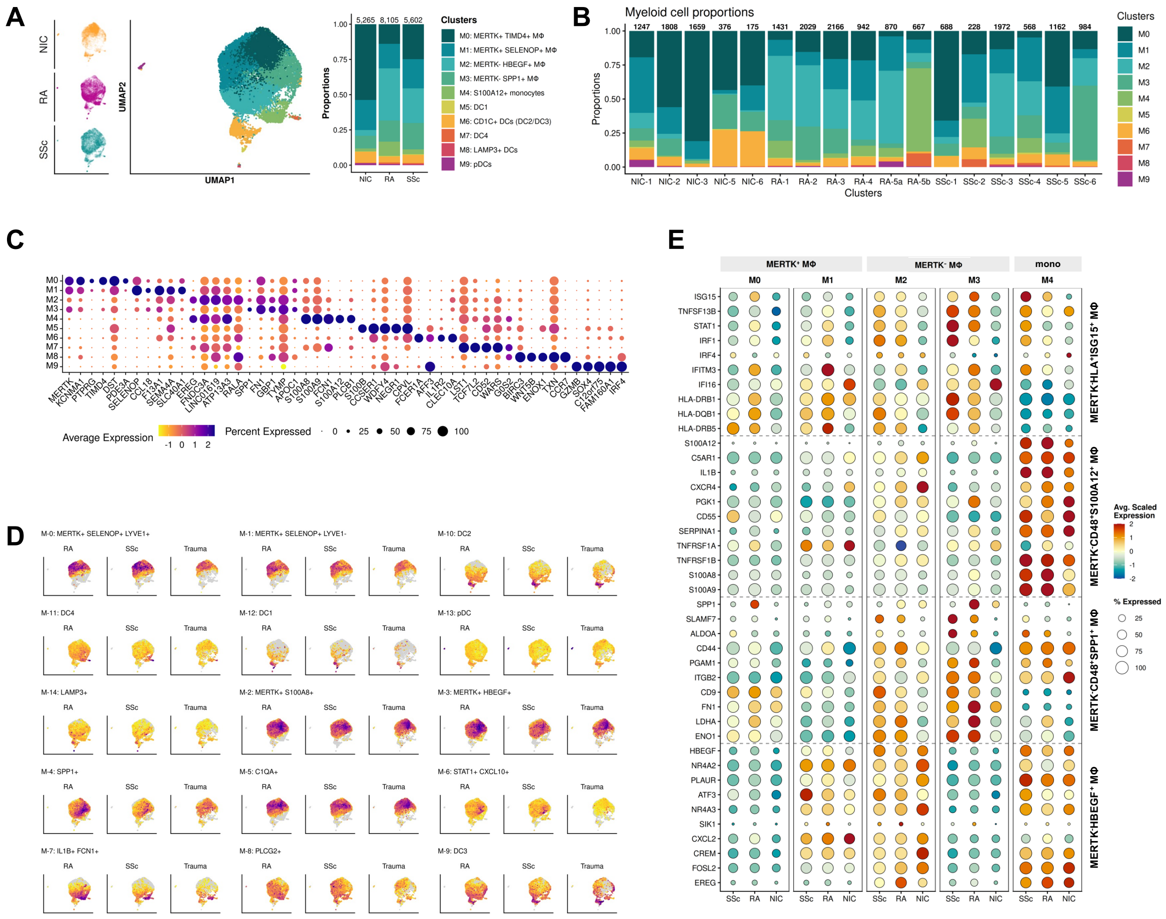


**Fig. S8. Characterization of myeloid cell populations.** (**A**) UMAP of re-integrated myeloid cell populations split by disease group. (**B**) Proportions of myeloid cell populations per biopsy; numbers above bars indicate cells per sample. (**C**) Top five marker genes per myeloid cell population. (**D**) Module scores of published myeloid cell subpopulation signatures (Zhang et al. 2023, M-0 to M-14) projected onto the myeloid cell UMAP, split by disease group (Trauma = NIC). (**E**) Marker genes of MERTK$^{-}$ populations in *(45)* (top three) and HBEGF macrophages in *(46)* (bottom).

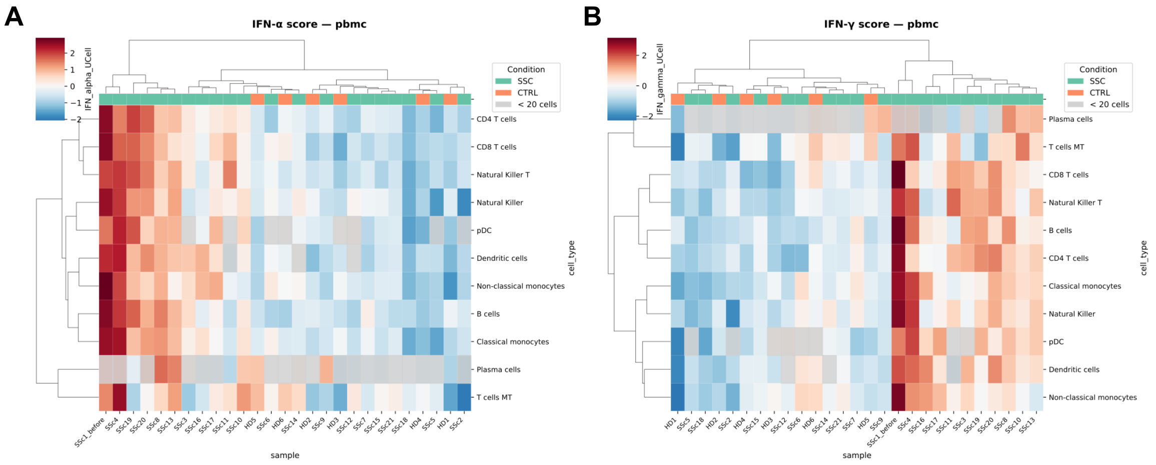


**Fig. S9. IFN response scores in circulating immune cells from SSc patients.** (**A**) Hallmark IFN-α and (**B**) IFN-γ Z-scores (pyUCell) per cell type and sample in a publicly available PBMC dataset (E-GEAD-872; Shimagami et al. 2025); Z-scores were computed per cell type relative to healthy controls (CTRL). Samples are hierarchically clustered (Euclidean distance, complete linkage).

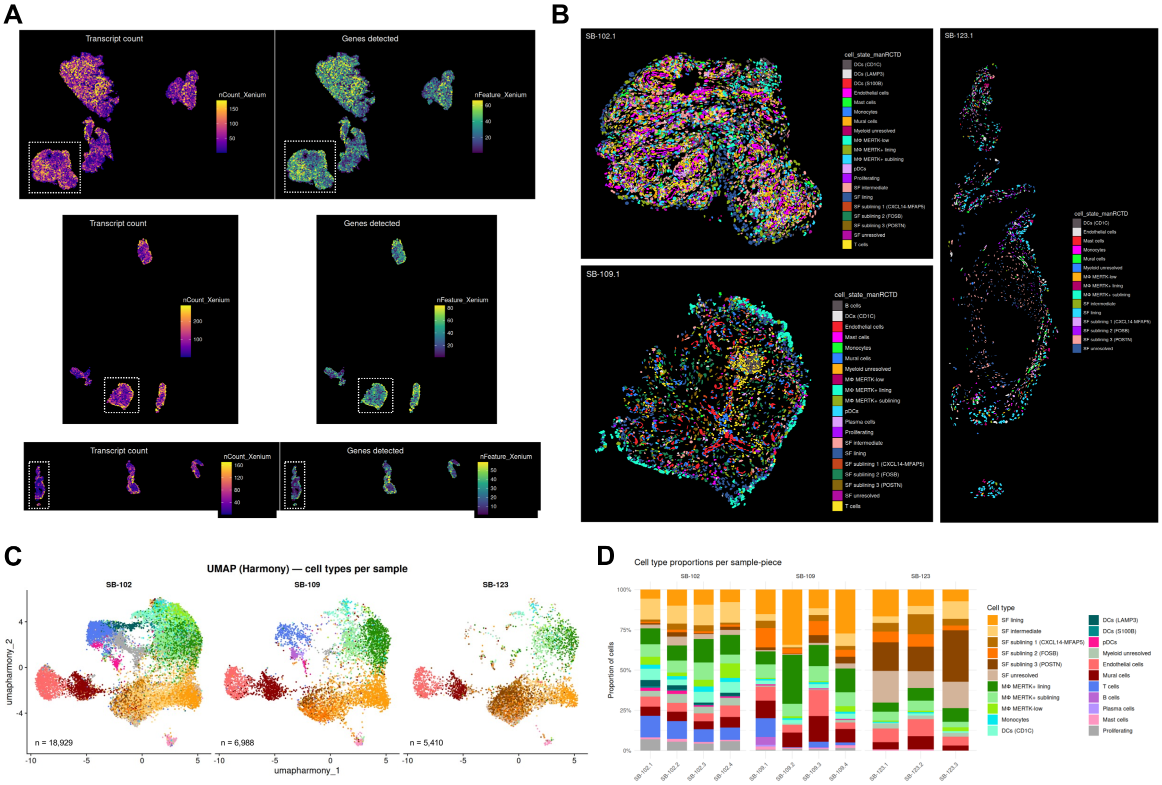


**Fig. S10. Xenium spatial transcriptomics annotation.** (**A**) Transcripts and gene detections per cell for each patient (top to bottom: SB-102, SB-109, SB-123); the tissue piece detailed in (B) is indicated. (**B**) Manual annotation of cell populations after clustering per patient and detailed RCTD subpopulation annotation of macrophages and SFs; cells with low marker gene expression were classified as "unresolved“. (**C**) Harmony-integrated UMAP across all patients with harmonized cell type labels and (**D**) cell type proportions per sample and tissue piece. For analyses shown in Fig. 5G–K, SF sublining subpopulations were merged into a single "SF sublining" category and DC subpopulations into "DCs."

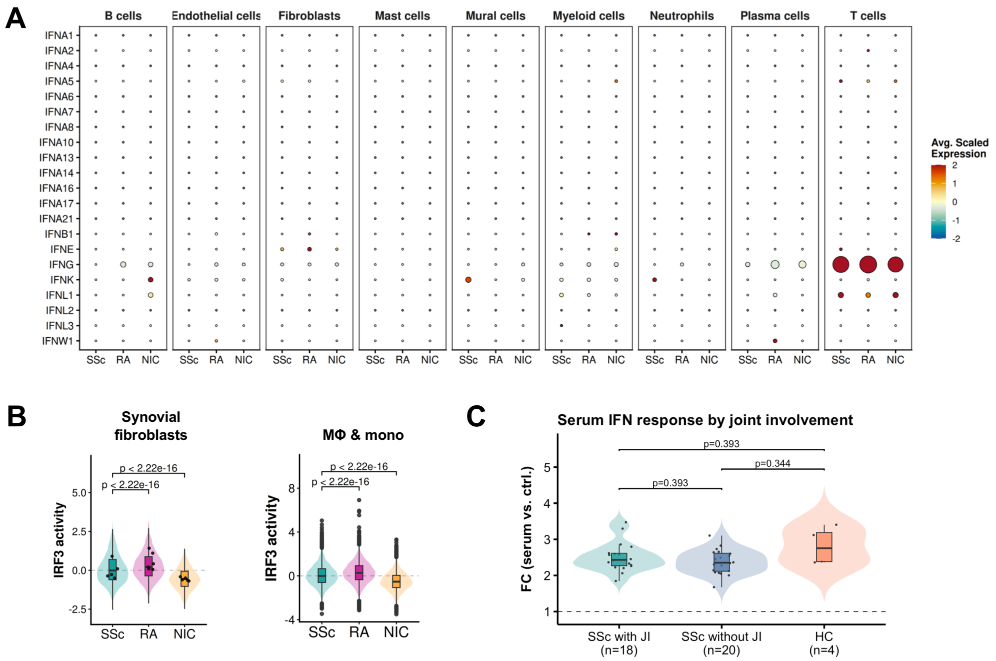


**Fig. S11. Assessment of local IFN production in SSc synovium and ISRE response.** (**A**) Expression of IFN transcripts across synovial cell types in SSc, RA, and NIC (dot size indicates percentage of expressing cells; color indicates scaled average expression). (**B**) Predicted IRF3 transcription factor activity in synovial fibroblasts and macrophage/monocyte populations across disease groups (decoupleR, CollecTRI; Wilcoxon rank-sum test). (**C**) ISRE-luciferase reporter activation in SF stimulated with sera from SSc patients (n = 38) grouped by joint involvement (JI) or healthy controls (HC, n = 4); Wilcoxon rank-sum test with BH correction.

**SUPPLEMENTARY TABLES**

Table S1: Pathohistology of full patient cohort

Table S2: Description and pathohistology for cohort subset used for scRNA-seq

Table S3: IFN-α2a serum level SSc cohort description

Table S4: ISRE-reporter serum stimulation SSc cohort description

Table S5: Custom 100-gene Xenium addon panel for spatial transcriptomics

Table S6: Antibody list and conditions in seqIF (Lunaphore COMET)
